# Mapping the brain-web morphology to characterise ageing and cognitive impairment

**DOI:** 10.64898/2026.11.06.764800

**Authors:** Francisco-Shu Kitaura, Aurelio Carnero Rosell, Juan Luis Sosa Ysalguez, Diego Anido Pereiro, Francesco Sinigaglia, Niels Janssen, Antonella Maselli, Ernesto Pereda, Marc Huertas-Company

## Abstract

Ageing reshapes human brain anatomy across spatial scales, yet its geometrical organisation remains difficult to quantify. Here we introduce `SHARP` (Spectral Hierarchy Analysis, Reconstruction and Prediction), which transfers hierarchical cosmic-web classification to structural MRI, identifying knots, filaments, sheets and voids at four spectral levels and combining their morphology with higher-order spatial statistics and anatomical measurements. In `CamCAN`, selected filament- and sheet-associated features show opposing correlations with age and distinct associations with subcortical and ventricular volumes. To quantify their contribution, we compare `SHARP` with a neural-network baseline combining 36 regional anatomical measurements, total brain volume and sex. Adding cosmologically motivated descriptors reduces the mean absolute error of MRI-only age prediction from 6.04 to 4.71 years (22%) in `CamCAN` and from 4.57 to 4.07 years (11%) in `OASIS-3` under participant-independent validation. Brain-web morphology with only brain volume and sex already achieves 5.12 years in `CamCAN`. For cognitive-status classification in `OASIS-3, SHARP` identifies 85% of Clinical Dementia Rating (CDR)-positive observations at 86% specificity, compared with 78% sensitivity for the anatomical baseline at matched empirical specificity. These results demonstrate that cosmological methods provide complementary geometrical information and measurable gains in characterising brain ageing and cognitive impairment.

## Main

Nature generates intricate web-like structures across remarkably different physical and biological systems, from the cellular cytoskeleton and branching biological networks to the large-scale distribution of matter in the Universe. Despite their different microscopic origins, these structures raise a fundamental question: whether common principles govern the emergence of complex spatial organisation across scales. A possible connection arises at the mesoscopic level, where collective geometrical patterns become apparent independently of the detailed dynamics of individual constituents. As recently explored in the context of complex structure formation [1], long-range interactions or information flow can introduce nonlocal coupling, allowing fluctuations to propagate across spatial scales and amplify initially small directional differences. Combined with anisotropic transport and nonlinear evolution, these processes can give rise to coherent sheets, filaments and interconnected concentrations, accompanied by higher-order correlations and departures from Gaussian statistics. In this perspective, the cosmic-web provides a particularly well-understood physical example of how complex geometrical organisation emerges from relatively simple initial conditions. Although gravity and biological processes operate through fundamentally different microscopic mechanisms, their resulting spatial structures may share geometrical and statistical properties that can be characterised within a common mathematical framework.

Supporting this perspective, Vazza and Feletti [2] identified similarities between the spatial power spectra of histological sections of human cerebral and cerebellar cortex and the matter distribution in cosmological simulations, after appropriate rescaling and normalisation. Their findings suggest that distinct physical systems can exhibit comparable statistical organisation despite their vastly different characteristic scales and underlying dynamics.

This motivates transferring hierarchical cosmic-web classifications developed in cosmology [3, 4] to structural brain imaging, not as a model of gravitational dynamics in the brain, but as a new geometrical language for identifying and quantifying its multiscale anatomical organisation. Combined with higher-order correlations [5], this approach provides complementary descriptions of spatial morphology and statistical relationships, opening a route to investigating how the organisation of brain tissue changes with ageing and cognitive impairment.

Ageing changes both the size of human brain anatomical regions and the organisation of tissue within them. Fluid-filled ventricles can expand, tissue can be lost, and the arrangement of grey and white matter can change [6–10]. These changes span many spatial scales. Brain-age models seek to bring this information together by learning how anatomy varies with chronological age. The difference between predicted and actual age, often called the brain-age gap, may help characterise departures from typical ageing, although its meaning for an individual’s health requires separate validation.

Anatomical atlases describe age-related changes through measurements of predefined brain regions. Complementary geometrical approaches can reveal information beyond regional volumes. Primus et al. 2025 [11] identified genetic associations with spectral descriptors of subcortical shape beyond those explained by volume, while

Escalante et al. 2025 [12] reported age-related patterns of global brain expansion and compression associated with cognitive performance and impairment. Cosmological power-spectrum and bispectrum statistics have also been used to predict brain age from MRI [5], although they do not explicitly classify local anisotropic morphology. Here, we introduce `SHARP` (Spectral Hierarchy Analysis, Reconstruction and Prediction)^1^, a framework that combines multiscale cosmic-web morphology with higher-order spatial statistics and conventional anatomical measurements.

Beyond brain-age estimation, we investigate whether these features capture structural differences associated with cognitive impairment. Rather than relying on the brain-age gap, we use the multidimensional `SHARP` representation directly to classify cognitive status, thereby assessing whether its morphological and statistical descriptors provide additional information beyond conventional anatomical measurements (Fig. 4). We define `BASE` as the reference model using 38 conventional inputs (total brain volume, 36 regional anatomical measurements and sex); `SHARP` adds 400 higher order statistics (HOS) and 128 spectral cosmic-web (BWEB) features, yielding 566 inputs.

Our aim is to track brain ageing through interpretable summaries that make structural changes easier to examine and assess. Such changes can be difficult to discern by eye in a three-dimensional MRI containing approximately 1.5 million brain-mask voxel intensities. Rather than relying exclusively on features learned directly from voxel intensities by neural networks, we construct a compact representation of mathematically defined descriptors with explicit geometrical and statistical interpretations. This allows us to examine the contributions of different spatial scales, morphological environments and higher-order correlations, while reducing the dimensionality of the subsequent prediction problem. One-dimensional curves of ordinary and reduced bispectra, together with histograms of cosmic-web morphology, provide a directly inspectable, quantitative description of how image structure varies across individuals and ages. These summaries could provide complementary quantitative descriptors for research and, subject to independent validation, clinical assessment. They may also help identify unusual patterns or possible acquisition and processing artefacts for further inspection. Their robustness to noise and acquisition differences remains to be established systematically.

### From cosmic structure to brain morphology

The search for a compact description of spatial geometry has a close counterpart in cosmology, where the large-scale distribution of matter is organised into the cosmic-web: an interconnected network of sheets, filaments and dense knots surrounding vast underdense regions known as voids. The importance of this analogy lies not in a similarity between the physical processes governing the Universe and the brain, but in the possibility of describing their spatial organisation using the same mathematical tools.

In the cosmological Zel’dovich picture, the cosmic-web emerges through anisotropic gravitational collapse, as matter contracts or expands at different rates along different directions [13]. This behaviour is characterised by the tidal tensor, *T*_*ij*_ = *∂*_*i*_*∂*_*j*_Φ, the Hessian of the gravitational potential Φ, where *i, j* ∈ {1, 2, 3} denote spatial coordinates. Its three eigenvalues describe the local tendency towards compression or expansion along three mutually perpendicular directions. Counting the directions of compression provides a natural geometrical classification: three, two or one correspond to knots, filaments or sheets, respectively, while the absence of compressive directions characterises voids. In practice, the classification is defined relative to an eigenvalue threshold [14–16]. This construction has been widely used to characterise cosmic structure and its connection to the distribution of dark matter haloes [17] (see Fig. 1).

**Fig. 1.**
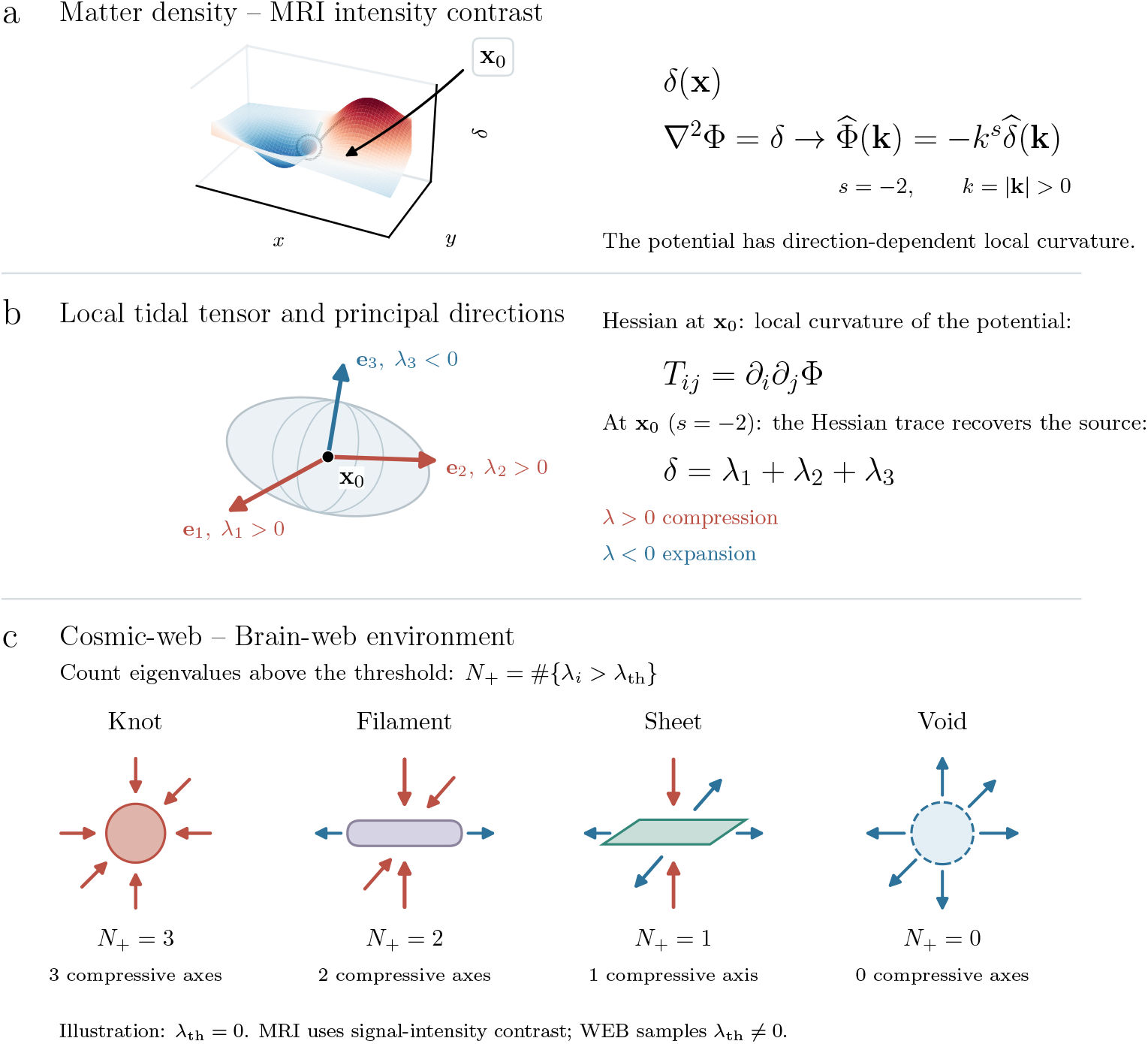
From a density field to brain-web morphology. **a**, Three-dimensional surface rendering of a two-dimensional slice through an illustrative matter-density contrast *δ*(**x**), with height indicating its amplitude. The marked point **x**_0_ identifies the location at which the associated gravitational potential is analysed in panel b. In cosmology, this contrast sources the gravitational potential through ∇^2^Φ = *δ*, with cosmological prefactors absorbed. The tidal tensor characterises the directional curvature of the potential, not of the density field itself. **b**, The tidal tensor *T*_*ij*_ = *∂*_*i*_*∂*_*j*_ Φ is the Hessian of the gravitational potential, evaluated locally at **x**_0_. Its eigenvectors identify three principal directions, and its eigenvalues describe compression or expansion in the cosmological tidal-web convention. **c**, The number of eigenvalues exceeding a threshold defines a knot (three), filament (two), sheet (one) or void (zero). The illustration adopts *λ*_th_ = 0; the analysis uses several positive and negative thresholds. For structural brain MRI, the matter-density contrast is replaced by the normalised MRI signal-intensity contrast. The resulting potential is a mathematical transform of image intensity, not a gravitational potential.

The connection to brain morphology becomes apparent when we consider how the gravitational potential is constructed. In Newtonian gravity, Poisson’s equation relates the potential to the matter density contrast *δ*. Absorbing the gravitational and cosmological prefactors into the potential gives

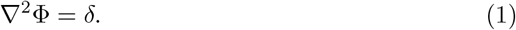

A three-dimensional field can be decomposed into Fourier modes with wavenumber *k* = 2*π/ℓ*, where *ℓ* denotes the corresponding wavelength. Small wavenumbers describe broad spatial variations, whereas large wavenumbers represent finer structure. Since the Laplacian corresponds to multiplication by −*k*^2^ in Fourier space,

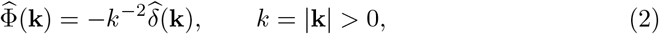

where hats denote Fourier-transformed fields. The gravitational construction therefore naturally introduces an inverse-square spectral weighting, emphasising large-scale fluctuations in the potential.

Although gravity has no direct dynamical role in the present description of brain anatomy, this construction can be transferred to MRI by replacing the matter density contrast with the normalised MRI intensity contrast. The resulting scalar field is not a gravitational potential, but its Hessian provides an analogous classification of local image geometry into knot-, filament-, sheet- and void-like environments. Importantly, these labels describe the curvature of a transformed intensity field, rather than distinct tissue types, individual neurons or neural connections.

This geometrical correspondence also suggests a natural extension. Hierarchical cosmic-web classification was previously introduced to capture non-local environmental information relevant to halo assembly bias [3]. Rather than restricting the classification to the gravitational *k*^−2^ weighting, we consider a hierarchy of spectral operators that probe complementary spatial scales, following Kitaura and Sinigaglia [4]. We define

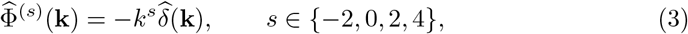

and apply the same Hessian-based classification to each transformed field. The index *s* specifies the spectral level: *s* = −2 recovers the gravitational construction, *s* = 0 preserves the relative amplitudes of the input frequencies in the scalar field, and *s* = 2 and *s* = 4 progressively emphasise finer spatial variations. The scale sensitivity of the final Hessian also includes its two spatial derivatives; it should not be identified with a narrow Fourier wavelength band. We refer to these levels as long-range, intermediate-range and short-range, respectively, although they represent relative scale sensitivities rather than sharply separated spatial intervals.

Hessian-based analysis is not new to medical imaging. Multiscale filters developed by Sato et al. 1998, 2000 [18, 19], Frangi et al. 1998 [20] and Lorenz et al. 1997 [21] use Hessian eigenvalues to identify and enhance tubular, sheet-like and blob-like structures, including applications to cerebral vessel segmentation [22]. These approaches typically construct the Hessian from a Gaussian-smoothed intensity field, with Fourier-space smoothing response exp(−*σ*^2^*k*^2^*/*2) and an additional second-derivative factor −*k*_*a*_*k*_*b*_. Apart from the overall sign convention, this resembles our *s* = 0 operator with additional Gaussian smoothing. The essential distinction is that conventional filters generally enhance selected structures through combinations of eigenvalue magnitudes and ratios, whereas our approach classifies the entire field into morphological environments according to the number of eigenvalues exceeding a threshold.

The four spectral levels therefore provide complementary geometrical descriptions of the same MRI rather than different images or anatomical segmentations. Figure 2 illustrates this hierarchy through axial and sagittal views of the input image and its derived morphological fields. The *s* = −2 classification captures broad spatial organisation, while the higher spectral levels increasingly reveal finer boundaries and texture. The scale dependence is also reflected in the normalised Fourier-space cross-correlation between the MRI intensity field and the derived morphology fields, which retains coherence out to progressively higher wavenumbers as *s* increases (see Methods, “Spectral brain-web construction”). Here, coherence refers to the correspondence between Fourier modes of the two fields, not merely agreement between their power-spectrum amplitudes. Although these correlations demonstrate that the morphological fields retain information about the spatial arrangement of image fluctuations, they do not establish independent predictive information. This is assessed separately through the feature-combination analysis. Moreover, the interpretation of high-wavenumber correlations must account for acquisition noise, preprocessing and sampling limitations.

**Fig. 2.**
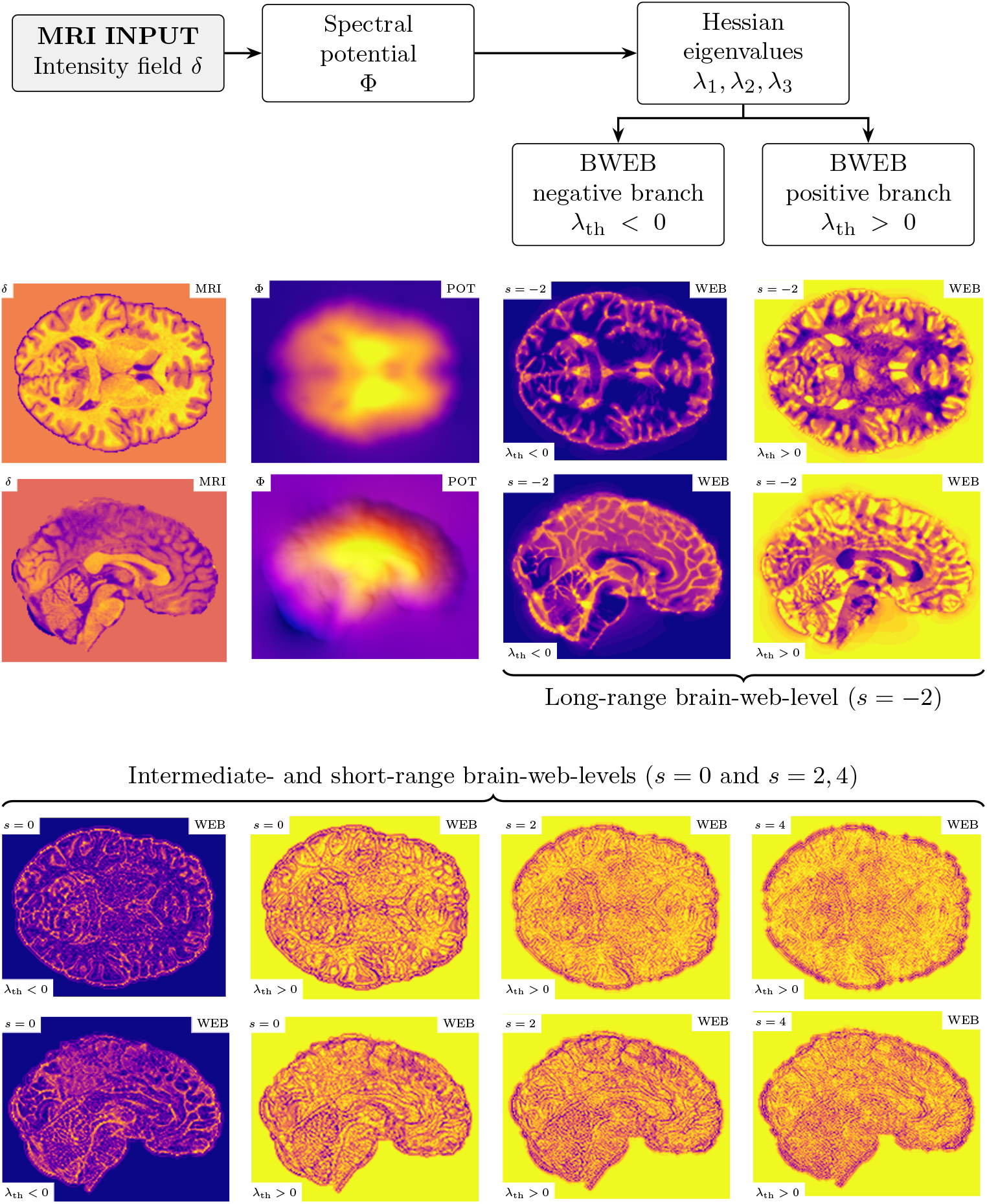
From MRI input to calculated spectral brain-web morphology. Axial (upper two rows) and sagittal (lower two rows) views of one subject from the `CamCAN` sample. Only the two panels labelled MRI show the input image intensity contrast, *δ* ≡ *I/*⟨*I*⟩_*M*_ − 1 within the brain mask *M*, with ⟨*I*⟩_*M*_ denoting the mean intensity in the mask and *δ* = 0 outside it; all panels labelled POT and BWEB are calculated from this input using our method. The first and third rows show, from left to right, the MRI input, its inverse-Laplacian scalar field Φ (POT), and the negative- and positive-threshold branches of the tidal-web classification associated with the *k*^*−*2^ kernel (*s* = −2). The second and fourth rows show the negative and positive branches for *s* = 0, followed by the positive branches for *s* = 2 and *s* = 4. These classifications are constructed from the Hessian eigenvalues of the spectrally weighted fields and provide complementary descriptions of image morphology, with higher spectral levels increasingly emphasising fine structure.

To obtain a compact representation that is less dependent on the choice of a particular classification boundary, we evaluate each spectral level over a range of eigenvalue thresholds, treating positive and negative thresholds separately. The numerical morphological labels are averaged at each voxel across the sampled thresholds, and the resulting scores are summarised in 16-bin histograms. Four spectral levels and two threshold-sign branches therefore yield eight histograms, collectively denoted BWEB (brain-web), containing 128 features. These distributions summarise threshold-averaged ordinal morphology scores, while discarding their precise voxel locations. The threshold sampling, numerical encoding and histogram construction are detailed in Methods.

Finally, we complement the BWEB descriptors with five bispectrum configurations, each containing 80 bins, which quantify higher-order spatial correlations. These include reduced bispectra normalised by combinations of power spectra, thereby incorporating two-point statistical information [5]. Together with total brain volume, sex and 36 regional anatomical measurements, the resulting `SHARP` representation contains 566 explicitly defined inputs, approximately 2,650 times fewer than the 1.5 million brain-mask voxel intensities of the analysed MRI (Fig. 3). This dimensionality reduction applies to the prediction inputs rather than the underlying image processing.

**Fig. 3.**
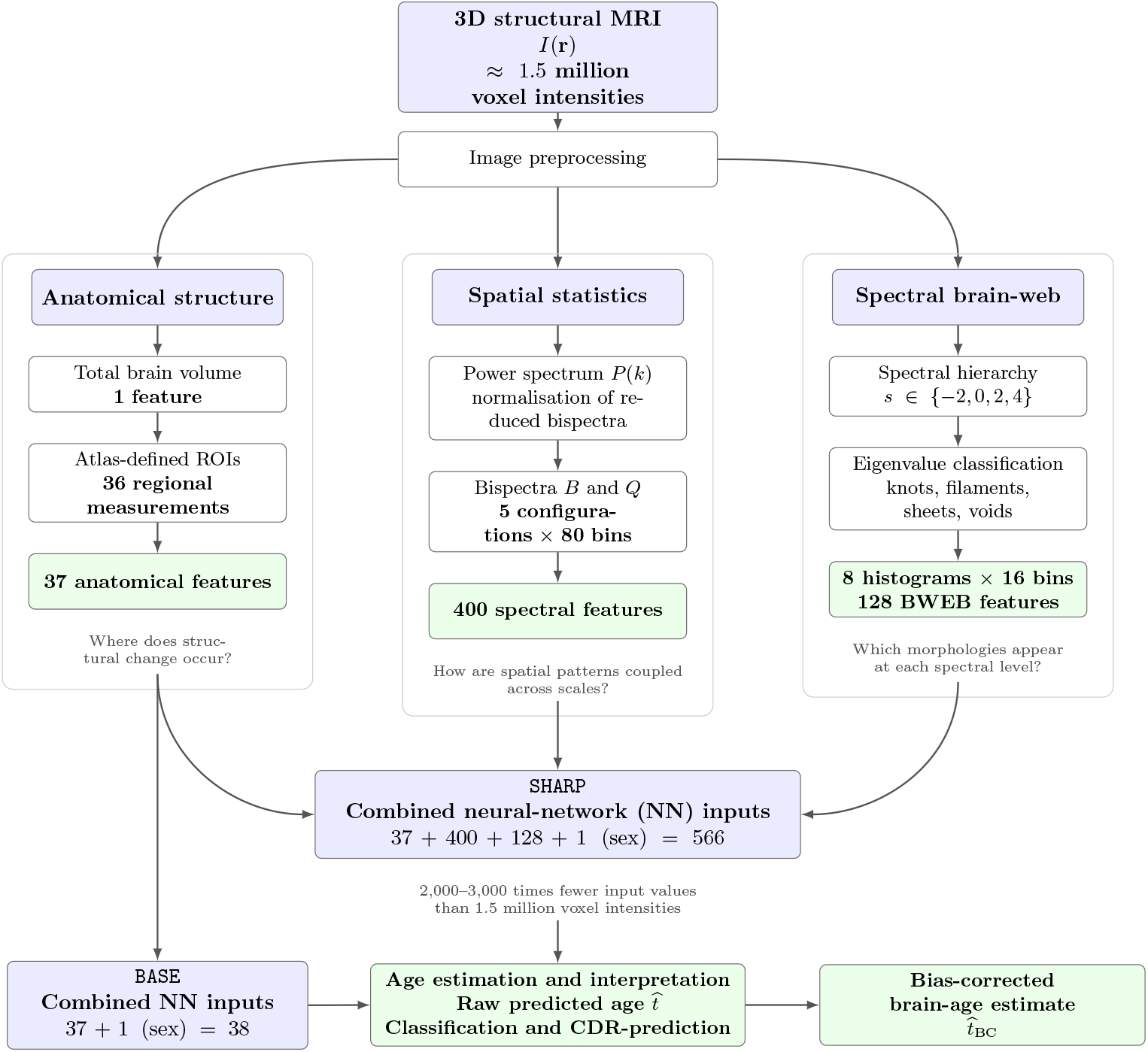
Schematic representation of the BASE and SHARP frameworks for brain-age prediction and cognitive-status classification. Each preprocessed structural MRI is characterised through three complementary branches. **Anatomy:** Total brain volume and 36 atlas-defined regional measurements provide 37 features describing anatomical size and structure. **Spatial statistics:** Five ordinary or reduced bispectrum configurations describe correlations among image patterns at different spatial scales. Each configuration is sampled in 80 bins, yielding 400 higher-order statistical (HOS) features. Reduced bispectra incorporate two-point information through normalisation by combinations of power spectra *P* (*k*). **Morphology:** Four spectral weightings, *k*^*s*^ with *s* ∈ {−2, 0, 2, 4}, probe complementary spatial scales. At each level, the eigenvalues of the Hessian classify local image geometry into knot-, filament-, sheet- and void-like environments. To reduce dependence on a single classification threshold, numerical morphological labels are averaged over multiple thresholds, separately for positive and negative branches. Each resulting field is summarised by a 16-bin histogram, producing eight histograms and 128 BWEB features. These distributions retain morphological information without preserving the spatial locations of individual voxels. **Prediction frameworks:** The baseline model (`BASE`) combines the 37 anatomical measurements with sex, yielding 38 neural-network inputs. Our proposed method (`SHARP`) additionally incorporates the 400 HOS and 128 BWEB features, yielding 37 + 400 + 128 + 1 = 566 inputs. `SHARP` therefore represents approximately 1.5 million brain-mask MRI voxel intensities through approximately 2,650 times fewer network inputs, while feature extraction still processes the original three-dimensional image. Both `BASE` and `SHARP` are evaluated for chronological-age prediction and cognitive-status classification. Age predictions can be subjected to an age-dependent bias correction, fitted independently of the test participants but requiring their known chronological age when applied. For Clinical Dementia Rating (CDR) classification, chronological age is included as an additional covariate, giving 39 inputs for `BASE` and 567 inputs for `SHARP`.

The resulting representation combines information about anatomical regions, spatial correlations and multiscale geometry, allowing their contributions to brain-age estimation and cognitive-status prediction to be evaluated together.

## Results: age estimation and cognitive-status prediction

We compare two feature-based approaches: the anatomical `BASE` model, which combines total brain volume, 36 atlas-derived regional measurements and sex (38 inputs), and `SHARP`, which additionally incorporates 400 higher-order statistical (HOS) features and 128 spectral brain-web morphological (BWEB) features (566 inputs; Fig. 3). For cognitive-status classification, chronological age is included as an additional covariate, giving 39 and 567 inputs, respectively. Clinical Dementia Rating (CDR) scores assigned by clinicians provide reference labels for both classifiers and are not predictors.

### Age estimation

To assess how much information these structural summaries carry about ageing, we tested how accurately they predict chronological age from an individual’s MRI scan. We compared combinations of morphological, volumetric and bispectrum features to examine whether they provide complementary information.

Age-regression performance was evaluated principally using subject-grouped cross-validation (SG), with all MRI sessions from a participant assigned to the same outer fold. We report uncorrected estimates (SG) and age-bias-corrected estimates (SGBC), with the correction fitted without using held-out participants (see Methods, “Subject-independent validation and age-related bias correction”). Unlike SG, SGBC requires the known chronological age of the evaluated participant and should therefore be interpreted as an age-adjusted brain-age estimate, not MRI-only age prediction. Figure 4c compares these results across feature combinations. For the full 566-input model, subject-grouped cross-validation (SG) yields a mean absolute error (MAE) of 4.07 years in `OASIS-3` and 4.71 years in `CamCAN`, with corresponding Pearson correlation coefficients of *r* = 0.81 and *r* = 0.94, respectively.

**Fig. 4.**
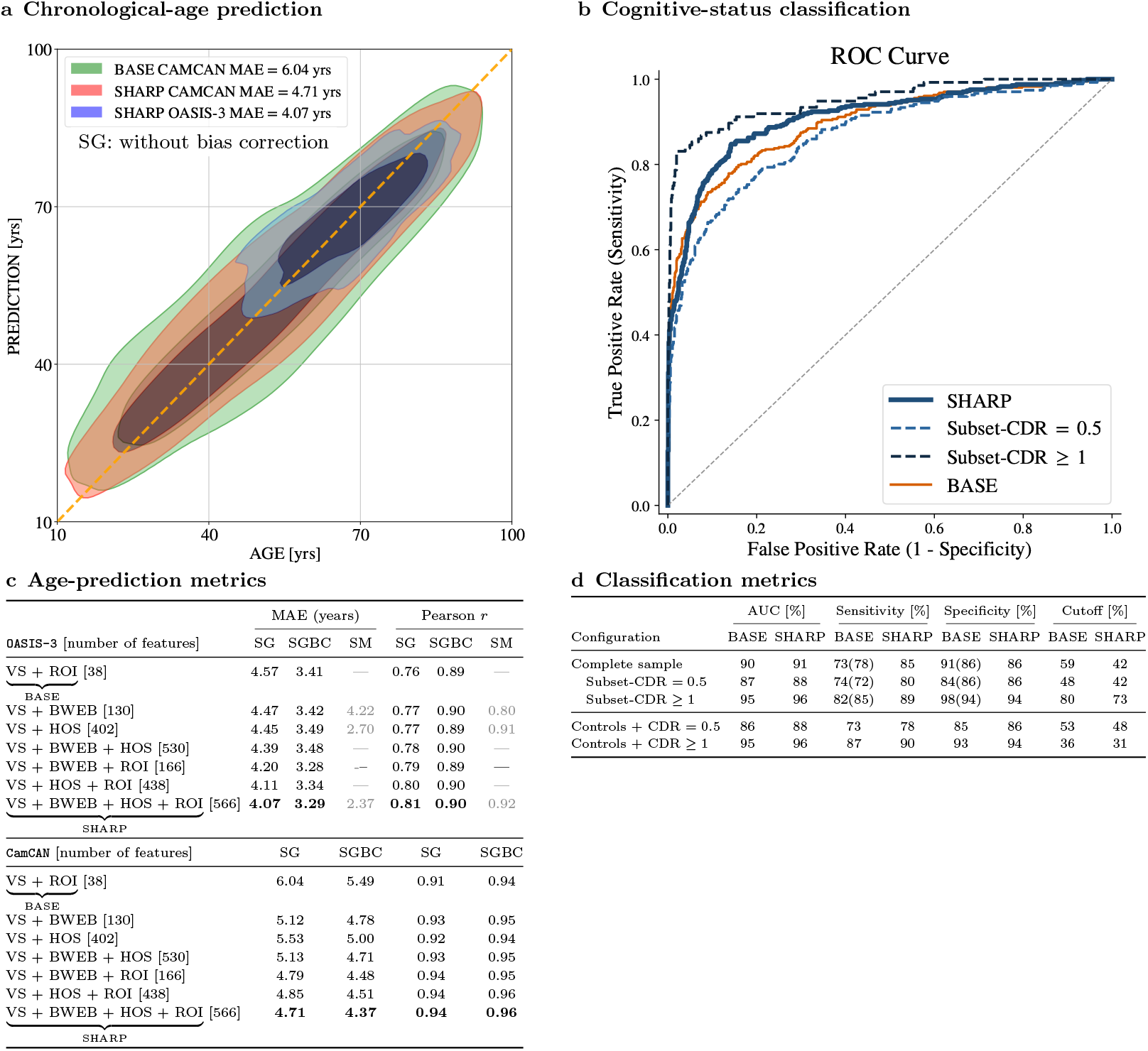
Quantitative performance of brain-age prediction and cognitive-status classification. **a**, Predicted versus chronological age for `OASIS-3` (blue), `CamCAN` for `BASE` (green) and `SHARP` (red), shown with one- and two-*σ* contours. **b**, Receiver operating characteristic (ROC) curves for distinguishing cognitively impaired (CDR-positive) observations from CDR-zero controls in OASIS-3, considering the complete sample and the CDR 0.5 and CDR ≥ 1 subgroups. **c**, Age-prediction performance for different feature combinations, quantified by mean absolute error (MAE) and Pearson correlation coefficient *r*. SG denotes subject-grouped cross-validation, SGBC includes age-dependent bias correction, and SM denotes session-mixed cross-validation. SG and SGBC use participant-disjoint folds, whereas SM may place sessions from the same participant in different folds. SGBC additionally uses known chronological age to correct the predictions. VS comprises total brain volume and sex; BWEB contains 128 morphological histogram features; HOS contains 400 bispectrum features; and ROI comprises 36 regional anatomical measurements. Numbers in brackets indicate the total feature count. `BASE` denotes VS + ROI (38 inputs) and `SHARP` denotes VS + BWEB + HOS + ROI (566 inputs). **d**, Cognitive-status classification performance for `BASE` and `SHARP` in terms of ROC area under the curve (AUC), sensitivity, specificity and probability cutoff. The indented rows evaluate CDR subgroups using the complete-sample classifier, whereas the final two rows correspond to subgroup-specific classifiers. Sensitivity, specificity and cutoffs are shown at the reported Youdenoptimal operating points, maximising *J* = Sensitivity + Specificity − 1. The operating cutoffs may differ between comparisons. Parenthesised entries in the `BASE` sensitivity and specificity columns are descriptive values obtained at the specificity of `SHARP` in the corresponding comparison (within rounding), not at the `BASE` cutoff listed in the table. Matched-specificity thresholds need not be identical numerical probability cutoffs.

Applying the age-dependent bias correction (SGBC) of Beheshti et al. 2019 [23] reduces the MAEs to 3.29 and 4.37 years, respectively, while increasing the corresponding correlations to *r* = 0.90 and *r* = 0.96.

These corrections reduce the reported MAEs by approximately 19% in `OASIS-3` and 7% in `CamCAN`, relative to the respective SG values. They remove part of the systematic dependence of the brain-age gap on chronological age, but because the Beheshti correction explicitly uses chronological age, SGBC measures age-adjusted agreement rather than MRI-only prediction accuracy. We therefore retain SG as the primary MRI-only generalisation metric and report SGBC alongside it, as is relevant to age-adjusted brain-age analyses [23–26].

In the auxiliary session-mixed (SM) analysis of `OASIS-3`, where repeated scans of the same participant may occur on both sides of a split, the full model achieves an MAE of 2.37 years and a Pearson correlation of 0.92. These values cannot estimate performance on previously unseen individuals. They nevertheless motivate prospective studies testing whether an earlier healthy scan can serve as a personalised anatomical baseline for later observations; the current SM experiment is not itself a longitudinal forecasting validation.

Interestingly, higher-order statistical (HOS) features show a much greater improvement under session-mixed validation, with the MAE decreasing from 4.45 years (SG) to 2.70 years (SM), compared with a modest decrease from 4.47 to 4.22 years for the BWEB morphological histograms. This contrast suggests that HOS may retain more participant-specific information than BWEB, although acquisition-related signatures and correlations between repeated scans could also contribute to the improvement.

Importantly, the `OASIS-3` age-prediction analysis reported by Carnero-Rosell et al. 2025 [5] used session-mixed cross-validation, which permits repeated MRI sessions from a participant to enter different folds. Those results therefore cannot quantify generalisation to previously unseen individuals. Re-evaluation of the corresponding volume, sex and higher-order statistical features (VS + HOS) under the present subject-grouped protocol gives an MAE of 4.45 years (SG), or 3.49 years after age-dependent bias correction (SGBC).

The feature-combination analysis in Fig. 4c shows a consistent overall trend across `OASIS-3` and `CamCAN`: complementing conventional anatomical measurements (`BASE`) with higher-order spatial statistics (HOS) and cosmic-web morphology (BWEB) improves brain-age prediction. The complete `SHARP` representation achieves the lowest uncorrected subject-grouped MAE in both cohorts. In `OASIS-3`, VS + BWEB + ROI gives a nearly identical SGBC MAE (3.28 years) compared with the complete model (3.29 years).

Relative to `BASE`, `SHARP` reduces the uncorrected, subject-grouped MAE by 0.50 years (approximately 11%) in `OASIS-3` and 1.33 years (approximately 22%) in `CamCAN`. The corresponding `BASE` SG MAEs are 4.57 and 6.04 years, respectively. These improvements refer to the MRI-only SG estimates, not the age-informed SGBC values. Figure 4a illustrates the `SHARP` age estimates after bias correction.

The correlations between cosmic-web morphology and chronological age reveal a pronounced dependence on spectral level (see Methods, “Anatomical associations across the hierarchy”). In `OASIS-3`, the strongest age associations are found at *s* = −2, corresponding to the classical cosmic-web classification based on the Hessian of the inverse-Laplacian potential. These correlations weaken at *s* = 0 and become nearly negligible at *s* = 2 and *s* = 4. Remarkably, `CamCAN` also exhibits strong age correlations at *s* = −2, but substantial correlations persist across the higher spectral levels (*s* = 0, 2, 4). This contrast suggests that broad morphological organisation captures robust age-related structural changes in both cohorts, whereas finer-scale morphological signatures are more clearly resolved in `CamCAN`.

The anatomical correlations of the BWEB descriptors reveal a hierarchy of morphological associations across spatial scales, particularly in `CamCAN`. At the classical tidal-web level, *s* = −2, the negative-threshold sheet-associated histogram bins correlate positively with age and ventricular and choroid plexus volumes, whereas filament-associated bins correlate positively with several subcortical volumes and negatively with age.

At the intermediate level, *s* = 0, both anatomical families remain represented, but their associations are redistributed among morphological environments depending on the eigenvalue-threshold sign. At the higher levels, *s* = 2 and *s* = 4, knot- and filament-associated bins in the negative-threshold branches show prominent correlations with subcortical, brainstem and cerebellar measurements, while sheet-associated bins correlate with ventricular and cerebrospinal-fluid measurements. The positive-threshold branches exhibit complementary associations.

These findings extend our previous bispectrum analysis [5], in which anatomical correlations differed between larger spatial wavelengths of approximately 17–33 mm, particularly associated with ventricular volumes, and finer wavelengths of approximately 5–8 mm, more sensitive to white-matter measurements. While the bispectrum probes explicitly selected Fourier wavenumbers, the BWEB hierarchy provides complementary geometrical classifications with progressively greater sensitivity to fine spatial variations.

For context, Barbano et al. 2026 [27] reported an uncorrected MAE of 3.96 years on `OASIS-3` using the adaptive-neighborhood method with training across multiple cohorts. Their proposed contrastive-learning method achieved 4.05 years under the same combined-cohort training strategy and 4.22 years when trained on `OASIS-3` alone.

For `CamCAN`, published structural-MRI MAEs include 5.33 years from Xifra-Porxas et al. 2021 [28] and 4.88 years from More et al. 2023 [29], the latter based on within-cohort evaluation of 650 participants and grey-matter, white-matter and cerebrospinal-fluid features.

Feng et al. 2020 [30] obtained an MAE of 4.21 years when evaluating on independent `CamCAN` data using a model trained on approximately 2,280 scans from a separate multisource sample.

A further reference is BrainAgeNeXt, a convolutional network trained on 11,574 MRI scans from 33 datasets, which reported a MAE of 2.78 years on its healthy-volunteer test sample [26]. In an independent evaluation of four pretrained brain-age models across different cohorts, Aguzin Parrilli and Belzunce 2026 [31] reported the lowest balanced global MAE for BrainAgeNeXt, 3.7 ± 0.2 years (mean and standard deviation across balanced subsamples). BrainAgeNeXt incorporates a Beheshti-type age-dependent bias correction. Training-set size, age distribution, preprocessing, bias correction and validation differences limit comparability; in particular, age-informed SGBC MAEs must not be presented as equivalent to raw MRI-only age-prediction MAEs.

### Cognitive-status prediction

We next examined whether the same multidimensional structural representation distinguishes CDR-positive observations (CDR *>* 0) from CDR-zero controls in `OASIS-3` (Fig. 4b,d). Rather than classifying participants from their brain-age gap, we trained dedicated classifiers on the `BASE` or `SHARP` features together with chronological age. All MRI sessions belonging to one participant were kept within the same cross-validation partition (see Methods, “Cognitive-status classifier”). We report ROC area under the curve (AUC), sensitivity and specificity at the stated Youden-optimal operating thresholds.

For the complete sample, `BASE` and `SHARP` achieve ROC AUCs of 0.90 and 0.91, respectively. At their respective Youden-optimal thresholds, sensitivity increases from 73% to 85%, while specificity changes from 91% to 86%.

To compare the methods at the same false-positive rate, we additionally evaluate `BASE` at the specificity achieved by `SHARP`. At 86% specificity, `SHARP` achieves 85% sensitivity compared with 78% for `BASE`. This advantage persists for CDR = 0.5 (80% versus 72% sensitivity at 86% specificity) and CDR ≥ 1 (89% versus 85% at 94% specificity).

We also trained classifiers separately to distinguish controls from CDR = 0.5 and CDR ≥ 1 observations. For CDR = 0.5, `SHARP` achieves an AUC of 0.88 compared with 0.86 for `BASE`, with sensitivities of 78% and 73%, respectively. For CDR ≥ 1, the corresponding AUCs are 0.96 and 0.95, with sensitivities of 90% and 87%. Full results, including specificities and operating thresholds, are reported in Fig. 4d.

## Discussion

Recent brain-age studies have demonstrated the predictive potential of structural MRI using regional anatomy, voxel-based methods and deep neural networks [26, 28– 30]. Complementary work on brain shape and spatial geometry has shown that anatomy carries information not fully represented by regional volume measurements [11, 12]. Here, `SHARP` investigates explicitly defined geometrical and statistical properties of structural MRI that carry information about ageing and cognitive status, complementing existing brain-age predictors. Its central novelty is the transfer of hierarchical cosmic-web classification to structural brain images, complemented by higher-order spatial statistics and anatomical measurements. Rather than learning image representations directly from voxel intensities, `SHARP` combines predefined BWEB morphological descriptors at four spectral levels and across both threshold-sign branches with higher-order statistical and anatomical features; the neural networks then learn predictive relationships from these descriptors. This also differs from conventional Hessian-based image filters, which generally enhance selected local structures [18, 20].

The resulting 566-feature representation has approximately 2,650 times fewer prediction inputs than the roughly 1.5 million brain-mask voxel intensities used in the analysis. This reduction applies to the prediction inputs; feature extraction still operates on the full MRI. Its advantage is thus primarily a compact and inspectable description of spatial structure, rather than an established reduction in computational cost or sensitivity to acquisition artefacts. Against the same anatomical baseline (`BASE`), `SHARP` reduces the uncorrected subject-grouped MAE from 4.57 to 4.07 years in `OASIS-3` and from 6.04 to 4.71 years in `CamCAN`, corresponding to improvements of approximately 11% and 22%, respectively. Notably, BWEB morphology together with only brain volume and sex already yields an MAE of 5.12 years in `CamCAN`, without regional anatomical or higher-order statistical features. The relative gains indicate that these morphological descriptors contain predictive information complementary to conventional regional anatomy, although they do not establish unique biological interpretations for individual features.

The cosmological connection provides a mathematical language for the directional structure. Its application to the brain, in turn, helped motivate a hierarchy extending beyond gravitational weighting. Interpretability here means that the image operations and summaries are explicitly defined. It does not mean that a knot or filament corresponds directly to a neural structure, or that a feature has a unique biological explanation. Establishing that connection requires comparison with anatomy, tissue properties and acquisition conditions.

The distinctive age associations of the classical tidal-web representation (*s* = −2) may be related to the mathematical properties of the inverse Laplacian in three dimensions. Its free-space Green’s function scales as *r*^−1^, corresponding to the inverse-square radial field implied by flux conservation for an isolated point source [32]. The Hessian of the resulting potential combines a local isotropic component, whose trace recovers the source field, with a nonlocal traceless component describing anisotropic spatial structure [33]. Applied to structural MRI, this decomposition provides a mathematical description of how local image-intensity contrast is related to the surrounding spatial organisation, without implying that brain anatomy is effectively governed by Newtonian gravitational dynamics or an inverse-square force law.

The scale-dependent BWEB associations provide a geometrical counterpart to the higher-order spatial correlations of brain ageing reported previously [5]. That bispectrum analysis related wavelengths of approximately *L* ~ 17–33 mm to ventricular measurements and broader anatomical changes, whereas shorter wavelengths (*L* ~ 5– 8 mm) were more sensitive to white-matter measurements. These wavelengths follow from *L* = 2*π/k* for the selected bispectrum configurations; they are not measured peak scales of individual BWEB environments. The BWEB hierarchy instead distinguishes geometric configurations across spectral weightings and threshold signs, without isolating a single Fourier wavelength.

A particularly notable feature of the classical *s* = −2 classification in `CamCAN` is the opposition between filament- and sheet-associated histogram bins in the negative-threshold branch: filament-associated features correlate positively with several subcortical volumes and negatively with age, whereas sheet-associated features correlate positively with age and ventricular and choroid plexus volumes. Although these anatomical changes are well established, their separation into eigenvalue-defined morphological environments supplies a geometrical description not explicitly represented by regional volumes or the bispectrum alone. At *s* = 0, 2, 4, associations with these anatomical families persist, but the dominant histogram bins shift across morphological environments and threshold branches. The prominence of *s* = −2 age associations in both cohorts, together with the stronger higher-*s* associations in `CamCAN`, suggests complementary sensitivities to broad and fine anatomical variation. The BWEB classifications themselves remain voxel-resolved at each spectral level and threshold, although their histogram summaries discard spatial location. This makes it possible to examine which anatomical regions contribute to these global associations in future spatially resolved analyses, without assuming that correlations with regional volumes already establish their anatomical origin.

The same geometrical and statistical representation also provides information relevant to cognitive status without reducing each MRI to a single brain-age gap. With chronological age included as a covariate for both classifiers, `SHARP` achieves an ROC AUC of 0.91 compared with 0.90 for `BASE`. At their respective reported Youden-optimal thresholds, sensitivities are 85% and 73%, with specificities of 86% and 91%, respectively. At matched empirical specificity of 86%, `SHARP` retains 85% sensitivity compared with 78% for `BASE`; for CDR = 0.5, the corresponding sensitivities are 80% and 72%. This operating-point advantage is particularly relevant when distinguishing CDR = 0.5 from CDR-zero observations, but the small difference in overall AUC does not imply a similarly large improvement across the ROC curve.

Discrepancies with clinical CDR labels are not, by themselves, evidence of changes preceding recognised impairment: model errors, acquisition effects and variability in clinical ratings remain possible explanations. Interpretation therefore requires independent clinical assessment rather than treating either prediction or recorded rating as infallible.

The reproducibility of clinical CDR assessments provides important context for interpreting our classification results. Previous studies have reported agreement with reference global CDR assessments of 83–87% [34, 35]. These findings indicate that clinician-assigned CDR scores, which constitute the reference labels for our models, are not free from assessment variability. Consequently, discrepancies between MRI-based predictions and clinical ratings may partly reflect uncertainty in the reference assessment rather than errors in the imaging-based classification alone.

Although these agreement rates concern the full CDR scale and cannot be interpreted as a strict upper bound on the accuracy of our binary classification tasks, they illustrate why near-perfect agreement with clinical labels should not be presumed. This is particularly relevant when distinguishing cognitively normal individuals from those with very mild impairment (CDR = 0.5), where clinical distinctions may be subtle.

Future studies using larger longitudinal cohorts, such as `ADNI` [36], could extend `SHARP` to ordinal classification across the full CDR scale, enabling comparisons with clinical assessments at different levels of impairment. Combined with longitudinal follow-up and independent clinical reassessment, such analyses could help distinguish prediction errors and variability in clinical ratings from structural changes preceding clinically recognised impairment.

Important limitations remain. The two cohorts were modelled separately, so cross-cohort transfer and robustness to new scanners or acquisition protocols have not been established. In particular, the weaker high-*s* age associations in `OASIS-3` may reflect acquisition heterogeneity, image quality, age sampling or other cohort differences; these possibilities cannot be separated by the present analysis. Independent-site validation, acquisition-controlled experiments and tests of feature stability will be needed before the proposed descriptors can support clinical use.

From a statistical perspective, the combination of isotropic two- and three-point correlations with anisotropic morphological descriptors moves towards a more complete characterisation of the multivariate distribution underlying structural MRI. A fuller description would require additional information, including higher-order correlations and spatial relationships between morphological environments.

The application of cosmic-web classification to structural MRI demonstrates that a geometrical framework developed in cosmology can also describe variations in human brain anatomy. This does not establish common mechanisms of structure formation in the two systems; rather, it supports the use of shared mathematical descriptors to investigate organisation across spatial scales. The approach provides a basis for further studies of how these morphological signatures relate to ageing, tissue properties and clinical outcomes [1].

## Ethics and reporting

This study used research data accessed under the relevant data-use procedures from the `OASIS-3` and `CamCAN` datasets. `OASIS-3` [37] is a de-identified neuroimaging and clinical research resource collected through the Washington University Knight ADRC. All procedures involving human participants in `OASIS-3` were approved by the Institutional Review Boards of Washington University in St. Louis, and written informed consent was obtained from all participants. All analyses were conducted in compliance with the `OASIS-3` Data Use Agreement, which prohibits attempts to re-identify participants and requires acknowledgment of the data source in all publications.

`CamCAN` [38, 39] is a population-based, multimodal (MRI, MEG, and cognitive-behavioural) dataset collected by the MRC Cognition and Brain Sciences Unit and University of Cambridge, comprising a cross-sectional adult lifespan sample. The data were collected in compliance with the Declaration of Helsinki and approved by the Cambridgeshire 2 Research Ethics Committee (reference: 10/H0308/50). Written informed consent was obtained from all participants.

The present study analysed de-identified data under the respective access agreements. No additional data collection involving human participants was performed by the authors.

The authors report approval from the Research Ethics and Animal Welfare Committee (CEIBA) at the University of La Laguna (CEIBA2024-3388; NJ).

## Data availability

`OASIS-3` is available through the `OASIS` data-access procedure [37]; `CamCAN` is available to approved researchers through its data-access portal [38, 39].

For reproducibility of `SHARP`, we intend to release the de-identified numerical features used in the regression and CDR-classification analyses, rather than the original MRI volumes. For each eligible subject or MRI session, the tabulated 566-input representation contains total brain volume and sex (two entries), 36 regional anatomical measurements, five 80-bin bispectrum configurations (400 entries), and eight 16-bin BWEB histograms (128 entries). The release will include the feature definitions, observation/participant grouping information and the analysis labels needed to reproduce the reported experiments, subject to the data-use and privacy requirements of the original cohorts. No raw or reconstructed MRI images will be redistributed. Public deposition of `OASIS-3`-derived participant/session level tables is contingent on verification of the dataset provider’s redistribution requirements.

## Code availability

The `SHARP` brain-age regression and cognitive-status classification software will be made available through the GitHub repository used by Carnero-Rosell et al. [5]: https://github.com/aureliocarnero/sharp.

## Acknowledgements

This research was primarily funded by the Cabildo de Tenerife through the IACTEC Technological Training Program, under the TF INNOVA grant. The funding was provided within the framework of MEDI-FDCAN 2016–2025, as part of the Tenerife Island Strategic Development Plan, in support of the `Cosmic Brain` project initiated in 2021 (Principal Investigator, PI: FSK). FSK also acknowledges the Spanish Ministry of Science and Innovation for financing the `Big Data of the Cosmic Web` project: PID2020-120612GB-I00/AEI/10.13039/501100011033 (PI: FSK), the *FIRE (Field-level bayesian Inference to Reconstruct the univErse)* project: PID2024-160504NB-I00 (co-PI: FSK), and the `Cosmology with Large Scale Structure Probes` project funded by the IAC (PI: FSK), which provided numerical tools and computing facilities. NJ acknowledges financial support from PID2021-127611NB-I00 (PI: NJ).

FSK and FS thank Kilian Peraza Vera for useful discussions on averaging cosmic-web regions for different eigenvalue thresholds.

The authors acknowledge the support of the IAC High-Performance Computing team and the computing facilities used in this research. They also acknowledge financial support for the publication fees from the CELESTE project (Grant Agreement No. 101136736), funded by the European Union under the WIDERA Teaming programme. Data were provided by `OASIS-3`: Longitudinal Multimodal Neuroimaging (Principal Investigators: T. Benzinger, D. Marcus, J. Morris; NIH P30 AG066444, P50 AG00561, P30 NS09857781, P01 AG026276, P01 AG003991, R01 AG043434, UL1 TR000448, R01 EB009352). AV-45 doses were provided by Avid Radiopharmaceuticals, a wholly owned subsidiary of Eli Lilly.

Data collection and sharing for this project was provided by the Cambridge Centre for Ageing and Neuroscience (`CamCAN`). `CamCAN` funding was provided by the UK Biotechnology and Biological Sciences Research Council (grant number BB/H008217/1), together with support from the UK Medical Research Council and University of Cambridge, UK.

## Author contributions

Francisco-Shu Kitaura conceived and led the study, proposed the application of cosmic-web morphology to structural brain imaging, and directed the development of the spectral hierarchy. He carried out the cosmic-web parameter studies for the `OASIS-3` and `CamCAN` datasets, contributed to software development, validation, data analysis and visualisation, secured funding, and led the writing of the manuscript.

Aurelio Carnero-Rosell developed and implemented the morphological analysis and cognitive-status classification pipelines. He applied the final analysis framework to both datasets, reviewed and refined the regression calculations, and conducted the final brain-age prediction analyses. He also contributed to software development, validation, visualisation and manuscript preparation.

Juan Luis Sosa Ysalguez investigated bispectrum configurations that improved predictive performance and performed the first joint brain-age regression analyses combining cosmic-web morphology and higher-order statistical features. This work formed part of his 2026 undergraduate dissertation in physics, supervised by Francisco-Shu Kitaura and Aurelio Carnero-Rosell.

Diego Anido Pereiro investigated the combination of two spectral levels of cosmic-web morphology for brain-age regression using methods other than neural networks, providing an early demonstration of the predictive potential of the hierarchical approach. This work formed part of his 2024 master’s dissertation, supervised by Francisco-Shu Kitaura and Aurelio Carnero-Rosell.

Francesco Sinigaglia contributed to the development of the cosmic-web spectral hierarchy and its associated methodological and numerical analyses.

Niels Janssen led MRI preprocessing for both cohorts and contributed to the cognitive-status classification analysis.

In addition to the above mentioned co-authors, Marc Huertas-Company, Antonella Maselli and Ernesto Pereda contributed to the methodological foundations of the study, including the development of higher-order statistical approaches and the broader analysis pipeline. They also participated in scientific discussions, interpretation of the results and preparation of the manuscript.

## Methods

### Study cohorts

We analysed T1-weighted structural magnetic resonance images from two independent cohorts, `OASIS-3` [37] and `CamCAN` [38, 39], using the same sample-selection criteria and preprocessing procedures as in Carnero-Rosell et al. [5]. We briefly describe both datasets below. Their age distributions and MRI power spectra are compared in Extended Data Figs. 1 and 2, respectively.

`OASIS-3` is a longitudinal neuroimaging study including cognitively normal adults and individuals at various stages of cognitive decline, whereas `CamCAN` provides a population-based cross-sectional sample spanning much of the adult lifespan. In both datasets, eligibility, clinical exclusion and image-quality criteria followed our previous analysis [5].

### OASIS-3

`OASIS-3` data were acquired through the Knight Alzheimer’s Disease Research Imaging Program at Washington University in St. Louis, Missouri, USA [37]. MRI acquisitions were obtained through three different Siemens scanner models (Siemens Medical Solutions USA, Inc.): Vision 1.5T, TIM Trio 3T and BioGraph mMR PET-MR 3T, with a heterogeneity of configurations.

The cognitively normal cohort comprised individuals classified as cognitively normal throughout available longitudinal follow-up; individuals who subsequently developed cognitive impairment were excluded. This sample contains 869 MRI sessions from 400 individuals aged 42–98 years (Extended Data Fig. 1). Chronological age was calculated at the MRI date with day-level precision.

For cognitive-status classification, we added 407 MRI observations from 336 individuals with positive CDR assessments. Of these observations, 271 had CDR = 0.5 (approximately 67%) and 136 had CDR ≥ 1 (approximately 33%). An MRI was assigned to the positive-CDR sample when it was acquired after a clinical assessment documenting a positive score. We use the CDR strata directly rather than equating CDR 0.5 with a clinical diagnosis of mild cognitive impairment.

#### CamCAN

Data used in this work were obtained from the `CamCAN` repository [38], based on the `CamCAN` study [39].

`CamCAN` is a population-based adult lifespan cohort [38, 39]. MRI data were acquired on a 3-T Siemens TIM Trio scanner using a 32-channel head coil. T1-weighted anatomical images were obtained with a three-dimensional MPRAGE sequence (TI/TR/TE = 900/2250/2.99 ms, flip angle 9°, field of view 256 × 240 × 192 mm, 1 mm isotropic voxels, GRAPPA factor 2; acquisition time 4 min 32 s). Age at scanning was recorded with year-level precision. The final cohort comprises 624 adults aged 18–88 years. The analysed sample is cross-sectional, with one T1-weighted MRI observation per participant; consequently, participant grouping does not alter the cross-validation partitions for `CamCAN`.

### MRI preprocessing

T1-weighted MRI data were processed following the pipeline used in our previous brain-age study [5]. Briefly, images were processed with version 4.3.0 of the HCP minimal processing pipeline [40], adapted when T2-weighted or field-map acquisitions were not available. Processing included spatial alignment, correction of image inhomogeneities and removal of non-brain tissue. Cortical reconstruction and volumetric segmentation were performed with FastSurfer [41–44], which provides FreeSurfer-compatible anatomical segmentations.

The scalar field used for the Fourier and BWEB analyses was the bias-field-corrected, skull-stripped and intensity-normalised T1-weighted image, rather than a tissue-probability or modulated tissue-volume map. Each image was represented on a common 300 × 300 × 300 Cartesian grid with 1 mm isotropic sampling, centred on the brain.

Let *I*(***x***) denote the processed MRI intensity and *M* the brain mask. The normalised intensity contrast used for the spectral analysis is

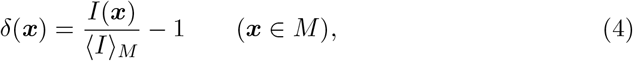

where ⟨*I*⟩ _*M*_ denotes the mean within the mask; the contrast is set to zero outside the mask.

### Spectral brain-web construction

The BWEB representation characterises each MRI through a hierarchy of four Hessian-based morphological classifications. Let 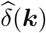 denote the Fourier transform of *δ*(***x***), with *k* = |***k***|. We define the spectrally weighted scalar fields

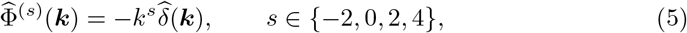

and construct at each level the symmetric Hessian tensor

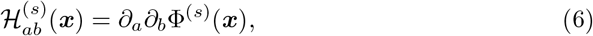

where *a, b* ∈ {1, 2, 3} denote the three spatial directions.

With the Fourier convention *∂*_*a*_ ↔ i*k*_*a*_, equations (5) and (6) give, for *k >* 0,

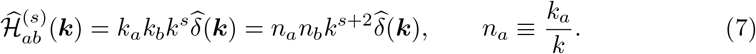

The factor *n*_*a*_*n*_*b*_ carries the directional dependence, whereas *k*^*s*+2^ determines the radial spectral weighting of the final tensor. At *s* = −2, this radial weighting is *k*^0^: the inverse-square factor belongs to the scalar potential, not to the final Hessian tensor. The four scalar fields Φ^(*s*)^ corresponding to *s* = −2, 0, 2, 4 are therefore ∇^−2^*δ*, −*δ*, ∇^2^*δ* and −(∇^2^)^2^*δ*, respectively. The *s* = −2 member reproduces the operator underlying the tidal-web construction, whereas increasing *s* progressively enhances high-frequency structure.

The zero Fourier mode was set to zero for the inverse-Laplacian (*s* = −2) calculation. This choice is immaterial for the Hessian because a spatial constant has vanishing second derivatives. Following the Fourier transformation, equation (7) was evaluated for all six independent components of the symmetric tensor and transformed back to real space. The resulting 3 × 3 tensor was diagonalised independently at each voxel.

For each MRI observation, we computed the normalised Fourier-space cross-correlation between the intensity contrast *δ* and the threshold-averaged BWEB morphology. The four environments were encoded as knots = 4, filaments = 3, sheets = 2 and voids = 1. For spectral level *s* and threshold-sign branch *η*, the threshold-averaged encoded field is 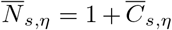, where 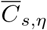 is defined below. We formed its dimensionless contrast within the mask,

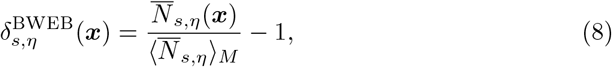

with zero contrast outside the mask, and evaluated

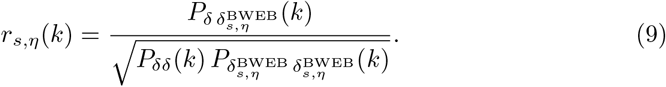

The numerator is the cross-power spectrum, and the denominator contains the two auto-power spectra. Spectra were averaged over modes within each wavenumber bin, with normalisation carried out for each observation before cohort or age-group averaging. The scale dependence of the resulting morphology can be examined through its Fourier-space cross-correlation with the MRI intensity field (Extended Data Figs. 3, 4 and 5). Across the sequence *s* ∈ {−2, 0, 2, 4}, the morphology fields retain higher correlations with the input image out to progressively higher wavenumbers, corresponding to smaller spatial scales through *ℓ* = 2*π/k*. This ordering connects the spectral construction to the image structures represented by each level: the lower levels primarily trace broad patterns, whereas the higher levels capture increasingly fine spatial variations.

A change in the correlation profiles occurs near the spherical Nyquist limit, *k*_Nyq_ = *π/*Δ*x* = *π* mm^−1^ for our Δ*x* = 1 mm grid. Below this limit, spherical Fourier shells are fully contained within the Cartesian sampling domain; above it, only a restricted set of directions remains available. Shell-averaged correlations beyond this scale therefore sample directions unevenly.

### Morphological classification and threshold averaging

Positive and negative threshold branches were sampled symmetrically about zero, excluding zero itself. For the *s* = −2 and *s* = 0 levels, threshold magnitudes were 0.01, 0.02, …, 0.09, followed by 0.10, 0.15, …, 0.30 (14 thresholds per branch). For *s* = 2, magnitudes ranged from 0.1 to 1.0 in steps of 0.1 (10 thresholds per branch), and for *s* = 4, from 2 to 10 in steps of 1 (9 thresholds per branch). The numerical classification labels were averaged voxel by voxel with equal weight across the sampled thresholds, separately for each spectral level and sign branch.

For both threshold-sign branches, voxels were classified by the number of eigen-values satisfying 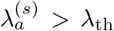: three, two, one or zero correspond to knots, filaments, sheets or voids, respectively. Increasing the positive threshold ultimately assigns every voxel to the void class, whereas decreasing the negative threshold ultimately assigns every voxel to the knot class. In either limit, the classification field becomes spatially uniform and contains no remaining morphological contrast.

Let 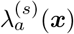, *a* = 1, 2, 3, be the eigenvalues of H^(*s*)^. For spectral level *s*, threshold-sign branch *η* ∈ {+, −} and threshold index *m*, let *C*_*s,η,m*_(***x***) ∈ {0, 1, 2, 3} denote the numerical morphological label assigned by the implemented eigenvalue criterion. Labels are averaged voxel by voxel:

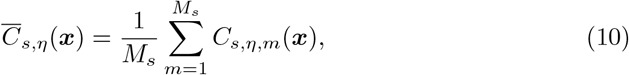

where *M*_*s*_ is the number of thresholds at spectral level *s*. The same ordered encoding is used at every threshold. Intermediate values are ordinal morphology scores: they summarise transitions between labels, but do not retain the complete class-membership distribution or constitute tissue probabilities.

For a histogram bin *ℬ*_*b*_, *b* = 1, …, 16, define the raw count

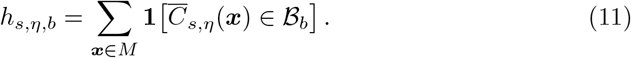

We denote the corresponding 16-bin feature vector by ***h***_*s,η*_ = (*h*_*s,η*,1_, …, *h*_*s,η*,16_); the bin boundaries are fixed across observations.

The concatenated representation is

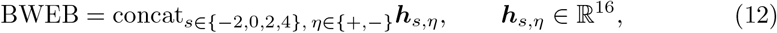

yielding 4 × 2 × 16 = 128 inputs. This operation differs from averaging four-component class-abundance histograms and from retaining the 4^4^ = 256 possible joint classifications across four levels.

Summaries of the histograms used in this study are shown in Extended Data Figs. 6, 7.

### Power-spectrum, bispectrum and regional features

Power-spectrum and bispectrum features were obtained from the same threedimensional MRI representation, following the methodology introduced in our previous analysis [5]. Fourier transforms were computed using FFTW. The images were represented on the 300^3^ mesh described above, and the MRI intensity field replaced the density field used in the original cosmological estimator. The numerical code was adapted from the estimator of Zhao et al. 21 [45], with physical spatial frequencies interpreted directly in mm^−1^. A Nearest-Grid-Point assignment scheme was used.

Bispectrum configurations were restricted to

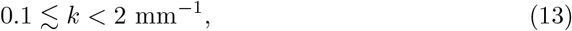

corresponding approximately to wavelengths from 3 to 63 mm. The lower cut reduces sensitivity to the largest-scale modes and image boundaries, whereas the upper limit restricts the analysis to frequencies supported by the sampled image field and reduces sensitivity to aliasing [46, 47].

The power spectrum describes the second-order Fourier statistics and the bispectrum the third-order statistics for closed wavevector triangles, ***k***_1_ + ***k***_2_ + ***k***_3_ = 0.

The ordinary bispectrum *B*_fluc_ is computed from the MRI intensity-fluctuation field *δ*, with *δ* = 0 outside the brain mask. The reduced bispectrum *Q* is instead computed directly from the MRI intensity field *I*:

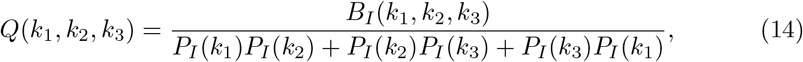

where *B*_*I*_ and *P*_*I*_ denote the bispectrum and power spectrum of the intensity field, respectively. Its normalisation therefore incorporates two-point statistics, expressing three-point correlations relative to the power at the participating spatial frequencies.

Following Carnero-Rosell et al. [5], each bispectrum configuration fixes two wavenumber magnitudes, *k*_1_ and *k*_2_, and samples the triangle angle *θ*. In the present analysis, each configuration is represented by 80 angular bins spanning 0 to *π*, with bin width Δ*θ* = *π/*80 = 2.25°.

To identify candidate bispectrum configurations for age prediction, we evaluated an ElasticNet model on the `CamCAN` dataset. We sampled the *k*_1_, *k*_2_ parameter space to identify the configurations yielding the lowest MAE. We then combined them iteratively until the MAE stopped improving. The selected configurations were subsequently evaluated on `OASIS-3`, where they improved upon the previously used configurations of Carnero-Rosell et al. 2025 [5], providing an additional cross-cohort check of the selected configurations. The selected representation contains three ordinary-bispectrum and two reduced-bispectrum configurations listed in Table 1.

**Table 1.** Bispectrum configurations selected using CamCAN and evaluated in both cohorts. Each configuration contributes 80 bins. Wavenumbers are in mm^*−*1^; the associated wavelength is 2*π/k*.

| Statistic | *k*<sub>1</sub> | *k*<sub>2</sub> |
|---|---|---|
| *Q* | 0.30 | 0.30 |
| *Q* | 0.10 | 0.15 |
| *B*<sub>fluc</sub> | 0.80 | 1.16 |
| *B*<sub>fluc</sub> | 0.15 | 0.26 |
| *B*<sub>fluc</sub> | 1.00 | 1.05 |

The regional anatomical features were derived from the FastSurfer segmentations [41–44]. FastSurfer provides FreeSurfer-compatible anatomical labels, allowing measurements to be associated with named brain regions. These region-based descriptors complement the BWEB histograms and bispectrum features by retaining an explicit anatomical localisation. Total brain volume was included separately as a global measure of brain size. The full model also retained 36 ROI measurements.

### Age-regression model

Chronological age was predicted from different combinations of global brain volume, sex, BWEB morphology, regional anatomical measurements and higher-order spectral statistics using a multi-input neural-network framework adapted from our previous study [5].

The complete `SHARP` regression representation combines 128 BWEB entries, 400 bispectrum entries, one total-volume measurement, one sex covariate and 36 regional measurements:

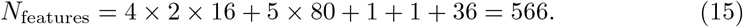

Chronological age is added for `SHARP` CDR classification, giving 567 inputs (39 for `BASE`). The ratio 1.5 × 10^6^*/*566 ≃ 2.65 × 10^3^ quantifies the input-dimensionality reduction, not a reduction in training sample size or model parameters.

Each feature family was processed through an independent fully connected network branch consisting of 128 and 64 neurons with leaky rectified linear unit activations. A dropout rate of 0.2 and batch normalisation were applied after the first dense layer. The resulting representations were concatenated and passed through shared dense layers of 128 and 64 neurons, with a dropout rate of 0.3 and batch normalisation after the first shared layer. The network terminated in a single linear output representing predicted chronological age. L1 and L2 kernel regularisation were applied to the dense layers, each with a coefficient of 0.001.

Training minimised the MAE using stochastic gradient descent with Nesterov momentum of 0.9 and an initial learning rate of 0.005, progressively reduced through a cosine-decay schedule. The schedule was defined over the maximum 300 training epochs, with a nominal minimum learning rate of 10^−4^ (a factor of 0.02 relative to the initial rate). Gradient norms were clipped at 1.0. The batch size was 32, with a maximum of 300 epochs. Early stopping monitored validation loss with a patience of 20 epochs, restoring the weights corresponding to the lowest validation loss. A new network, optimiser, cosine-decay schedule and batch-normalisation state were initialised for every outer fold.

For each training fold, the input features were standardised by subtracting their training-set means and dividing by their training-set standard deviations. The resulting transformations were subsequently applied to validation and test observations without refitting. Missing and non-finite feature values were replaced by zero before standardisation.

### Subject-independent validation and age-related bias correction

For the primary age-regression analyses, we used ten-fold stratified group cross-validation. Chronological ages were divided into ten quantile-based bins (with duplicate bin boundaries removed where necessary), while participant identifiers defined the groups. All MRI sessions of a participant were assigned to the same outer fold; thus, outer test participants were excluded from network training, early stopping and bias calibration.

Within each outer development set, a grouped random split reserved approximately 10% of the participants for internal validation. A second group-based split of the validation participants formed two non-overlapping subsets: one for early stopping and the other for age-bias calibration. All four subsets (network training, early stopping, calibration and outer testing) were therefore participant-disjoint.

We denote the uncorrected held-out age predictions as SG and the corresponding Beheshti-corrected estimates as SGBC. This notation is retained for both cohorts for consistency: grouping is necessary for repeated observations in `OASIS-3`, whereas for the one-scan-per-participant `CamCAN` sample the corresponding folds are participant-independent without an additional grouping constraint. The auxiliary OASIS-3 session-mixed (SM) analysis may place repeated scans of a participant in different outer folds and therefore does not measure generalisation to unseen participants.

#### Age-related bias correction

Brain-age prediction models commonly exhibit systematic age-dependent bias, tending to overestimate chronological age in younger individuals and underestimate it in older individuals. Several correction procedures have been proposed to address this effect [23–25]. Recent independent benchmarking has further highlighted the importance of age-related bias and validation procedures when assessing the generalisability of brain-age models [31].

We investigated different correction approaches, including the residual-based method proposed by Beheshti et al. 2019 [23], inverse affine correction and MAE-optimised affine calibration. The residual-based correction of Beheshti et al. provided the lowest MAE in our experiments and was therefore adopted for the final analysis.

Specifically, we define the uncorrected brain-age difference as

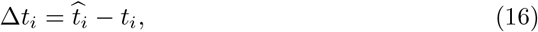

where 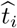 and *t*_*i*_ are the predicted and chronological ages of observation *i*, respectively.

Following Beheshti et al., we model the systematic dependence of this difference on chronological age through a linear relation,

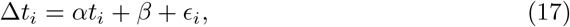

where *α* and *β* are the fitted slope and intercept and *ϵ*_*i*_ denotes the residual. The coefficients were estimated separately within each outer fold by ordinary least-squares regression of the calibration-subset residuals 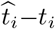 against chronological age *t*_*i*_. The calibration observations were not used to fit network weights or select the early-stopping epoch.

The bias-corrected brain-age estimate is then obtained by subtracting the fitted age-dependent offset:

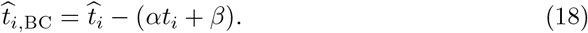

The fitted residual correction was evaluated at the known chronological age of each held-out test observation and subtracted from its uncorrected network prediction. The coefficients *α* and *β* were estimated solely from the internal calibration observations, without using the outer test fold. The early-stopping and calibration subsets were separated by participant under the revised subject-grouped validation scheme.

Importantly, although the fitted coefficients are independent of the held-out observations, the correction itself explicitly uses their chronological ages. As discussed by de Lange and Cole [25], this procedure reduces age-dependent prediction bias and can substantially reduce MAE, but the resulting corrected estimate should not be interpreted as an MRI-only prediction of chronological age.

We therefore report both the original subject-grouped predictions (SG) and the corresponding bias-corrected estimates (SGBC), allowing their performance to be distinguished. The code also evaluates two alternative calibrations on the same held-out network predictions: the inverse-slope (Cole-type) correction, 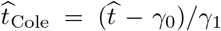, whose coefficients are fitted from predicted versus chronological age in the calibration subset, and a directly MAE-optimised affine transformation 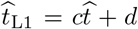. The SGBC results reported here correspond to the Beheshti residual correction. The choice among correction methods was examined separately from the uncorrected SG metric, which remains the primary evaluation. MAE and Pearson *r* were calculated separately for the held-out SG and SGBC estimates. Because SGBC uses chronological age at evaluation, an increase in its correlation does not demonstrate improved MRI-only age-predictive information.

### Anatomical associations across the hierarchy

We examined two complementary relationships: the spatial correspondence between MRI intensity and the threshold-averaged morphology fields, and the associations of BWEB histogram bins with regional anatomy and chronological age. The image-field cross-correlations are similar across the age groups shown, with comparatively narrow uncertainty bands in `OASIS-3` and `CamCAN` (Extended Data Figs. 3–5). This consistency supports a reproducible relationship between image intensity and the derived morphology within the analysed samples. It does not imply that the morphology is independent of age: stable overall profiles can coexist with systematic age-related changes in individual features.

To investigate the anatomical context of the BWEB morphological descriptors, we examined their associations with conventional morphometric measurements and age, complementing earlier analyses based on statistical descriptors of brain structure [5]. The *s* = 2 and *s* = 4 levels showed closely related correlation profiles, suggesting overlapping information, whereas the *s* = −2 level showed more distinct associations. In `OASIS-3`, associations with age weakened at *s* = 0 and became nearly negligible at *s* = 2 and *s* = 4 (Extended Data Figs. 8 and 9). This suggests that the higher-level BWEB histograms capture less age-related information in this cohort. It does not imply that all fine-scale information is uninformative: bispectrum features still improved prediction.

In `CamCAN`, the histogram bins showed clear positive and negative associations with age across the spectral hierarchy, with patterns depending on spectral level and threshold sign (Extended Data Figs. 10 and 11).

At *s* = −2 and negative thresholds (*λ*_th_ *<* 0), bins associated with sheet-like morphology correlated positively with age, lateral and third ventricle volumes, and choroid plexus volume. Filament-associated bins correlated positively with several subcortical volumes and negatively with age. Recurrent associations with ventricular and subcortical structures provided qualitative agreement between cohorts, although their strengths differed. These across-subject associations connect the descriptors to age-related anatomy and are distinct from the stable image-field cross-correlations described above. Shared correlations with age do not, by themselves, identify which anatomical changes cause a particular BWEB response.

The absolute errors also reflect differences in age sampling. `CamCAN` spans a broader age range than `OASIS-3`, which is concentrated at older ages. Using approximate chronological-age standard deviations of 10.0 years for `OASIS-3` and 18.4 years for `CamCAN`, the full-model SG MAEs correspond to approximately 41% and 26% of the respective age dispersions, while the SGBC MAEs correspond to approximately 33% and 24%. Since SGBC uses chronological age, these normalised quantities are descriptive rather than independent measures of MRI-only prediction accuracy. Models were trained separately within each cohort, without evaluating cross-cohort transfer.

Acquisition differences offer a possible explanation. `OASIS-3` combines multiple scanner models and protocols, including 1.5-T and 3-T acquisitions with heterogeneous acquisition times, whereas `CamCAN` used a consistent 3-T protocol with a uniform acquisition time. These differences can affect image contrast, noise and effective resolution even when nominal voxel sizes are similar. The power-spectrum ratio decreases from approximately 1.5–1.6 at large scales to about 1.15 at the smallest plotted scales. After normalising by the large-scale amplitude ratio, this corresponds to a relative small-scale power deficit of approximately 25% in `OASIS-3` compared with `CamCAN` (Extended Data Fig. 2). This suppression of fine-scale power, together with its greater variability, may contribute to the weaker age sensitivity of the higher spectral brain-web levels in `OASIS-3`. The observations are consistent with acquisition-related effects, although they do not isolate these from biological variability or preprocessing differences.

### Cognitive-status classifier

The regression network was adapted for binary CDR classification by retaining its feature branches, shared layers, regularisation and optimiser schedule and replacing the linear output with a single sigmoid unit.

Chronological age was concatenated as an additional predictor. For binary label *y*_*n*_ ∈ {0, 1} and predicted sigmoid score *p*_*n*_, the supplied production code uses weighted binary cross-entropy by default,

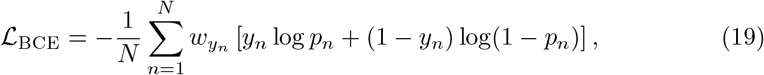

where the weights are *w*_*c*_ = *N/*(2*N*_*c*_) for class *c*, with *N*_*c*_ estimated in each training fold. This expression reduces to ordinary binary cross-entropy when weighting is disabled. Both binary accuracy and ROC AUC were tracked during training. In the supplied production code, early stopping monitors validation AUC by default, with a patience of 20 epochs. ROC curves and AUC were used to report discrimination (Fig. 4b,d). A sigmoid output is not by itself evidence of probability calibration, especially with class-weighted training.

In the `BASE` configuration the classifier receives 38 morphometric and demographic inputs, including sex, plus chronological age (39 total), whereas `SHARP` additionally receives 400 HOS and 128 BWEB inputs (567 total). Each uses the same clinician-assigned CDR labels for supervised training. Three training configurations were considered: controls versus all positive-CDR observations; controls versus CDR 0.5; and controls versus CDR ≥ 1. For the combined model, performance was also evaluated separately for each positive-CDR subgroup against controls. These subgroup evaluations use the combined model’s predictions, whereas the last two rows of Fig. 4d correspond to separately trained models.

For CDR classification, outer cross-validation splits were stratified by CDR category at the participant level, with every MRI session of a participant assigned to the same fold. Participant-disjoint internal validation sets were used for model selection. Where repeated ensemble fits were used, predictions were combined per held-out MRI session before calculating ROC AUC and operating-point statistics. ROC AUC was evaluated using held-out participants. Sensitivity and specificity are reported at the Youden-optimal operating points of the ROC summaries, with *J* = Sensitivity + Specificity − 1.

The CDR classifiers reported here use the morphometric, HOS and BWEB representations (as applicable), with chronological age as an additional covariate; they do not use predicted brain age, the brain-age gap or an age-bias-corrected brain-age estimate as classifier inputs. The SG/SGBC terminology is reserved for the regression results. Incorporating a corrected brain-age feature in a future classifier would require independently trained, participant-disjoint regression and calibration stages.

Clinical Dementia Rating (CDR) is a clinical staging assessment rather than an independent pathological diagnosis [48]; we use clinician-assigned scores as reference labels, not as imaging-derived inputs.

### Cross-validation and statistical evaluation

All principal age-regression and cognitive-status performance metrics were calculated from subject-grouped held-out predictions. Sessions of a participant never appeared in both the outer training and test folds. Session-mixed OASIS-3 results are shown only as descriptive comparisons and do not support claims of independent-subject generalisation. The age-regression SGBC metrics use the correction in Eq. (18); SG metrics use the raw network outputs. MAE and Pearson *r* are calculated separately for the corrected and uncorrected held-out estimates. The reported CDR classification does not include a corrected brain-age input; age correction is applied only to the separate regression analysis.

Brain-age prediction was evaluated using 1,000 randomised realisations of ten-fold subject-grouped cross-validation. In each realisation, all MRI sessions from a given participant were assigned to the same outer fold, ensuring that no scans from a test participant contributed to model training, early stopping or bias calibration. The training, early-stopping and calibration subsets were also separated by participant, and feature standardisation was fitted exclusively on the training data.

Each MRI observation received one out-of-fold prediction per realisation. The final uncorrected brain-age estimate was obtained by averaging the 1,000 predictions for each observation:

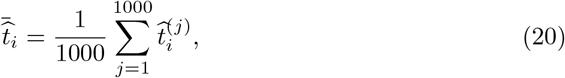

where 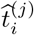 denotes the held-out prediction for observation *i* in realisation *j*.

The reported MAE and Pearson correlation were calculated from these ensemble-averaged predictions, rather than by averaging the performance metrics of the individual realisations. The raw predictions constitute our primary MRI-only results (SG). Beheshti-type age-bias correction was performed separately (SGBC), using calibration coefficients estimated without the test participants but requiring their known chronological ages when applied.

For *N* observations with chronological ages *a*_*n*_ and predicted ages *a*_*n*_, the mean absolute error was

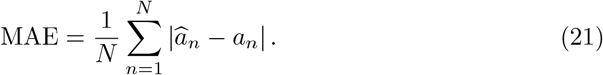

Pearson’s correlation coefficient between observed and predicted ages was

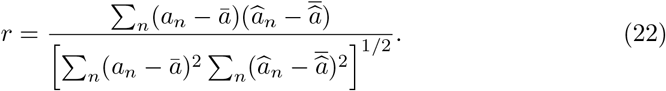

### Morphological and image-field associations

Associations between BWEB histogram bins, chronological age and regional anatomical measurements follow the morphometric analysis of Carnero-Rosell et al.[5]. The anatomical heatmaps report Kendall rank correlations separately for each spectral level and threshold-sign branch. In `OASIS-3`, associations between BWEB histogram bins and age weaken at higher spectral levels and become nearly negligible at *s* = 2 and *s* = 4 (Extended Data Figs. 8 and 9). In `CamCAN`, the bins show clear positive and negative associations with age across the spectral hierarchy (Extended Data Figs. 10 and 11). This contrast indicates weaker age sensitivity of the fine-scale BWEB summaries in `OASIS-3`.

**Extended Data Fig. 1.**
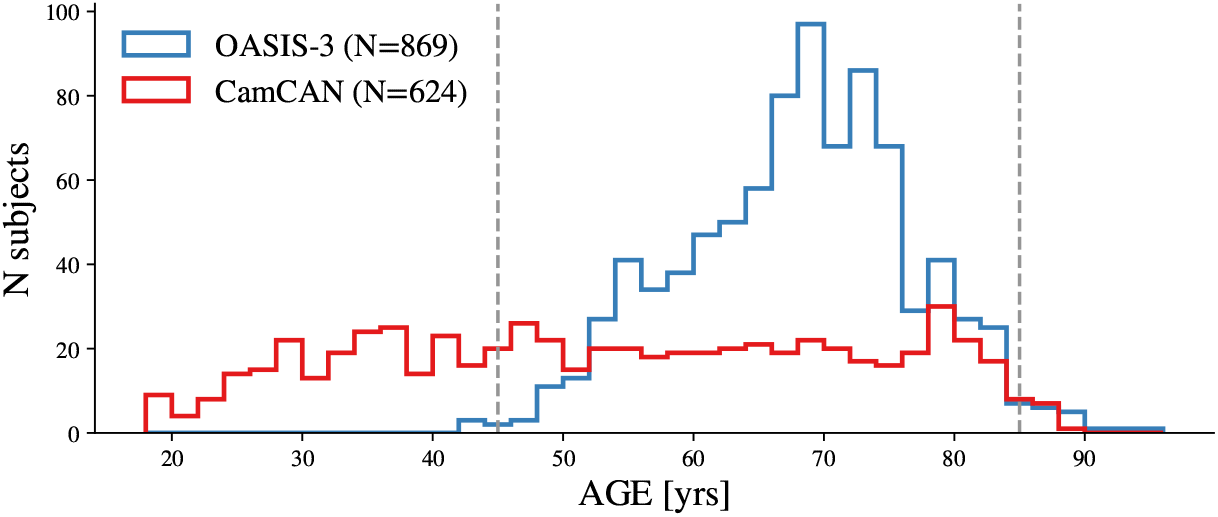
Age distributions of the `OASIS-3` (blue; *N* = 869) and `CamCAN` (red; *N* = 624) samples used in this study. Histograms show the number of MRI observations in each age bin. `OASIS-3` is concentrated at older ages, with denser sampling over a narrower age interval, whereas `CamCAN` spans a broader and more evenly sampled age range. These differences should be considered when comparing absolute age-prediction errors between cohorts.

**Extended Data Fig. 2.**
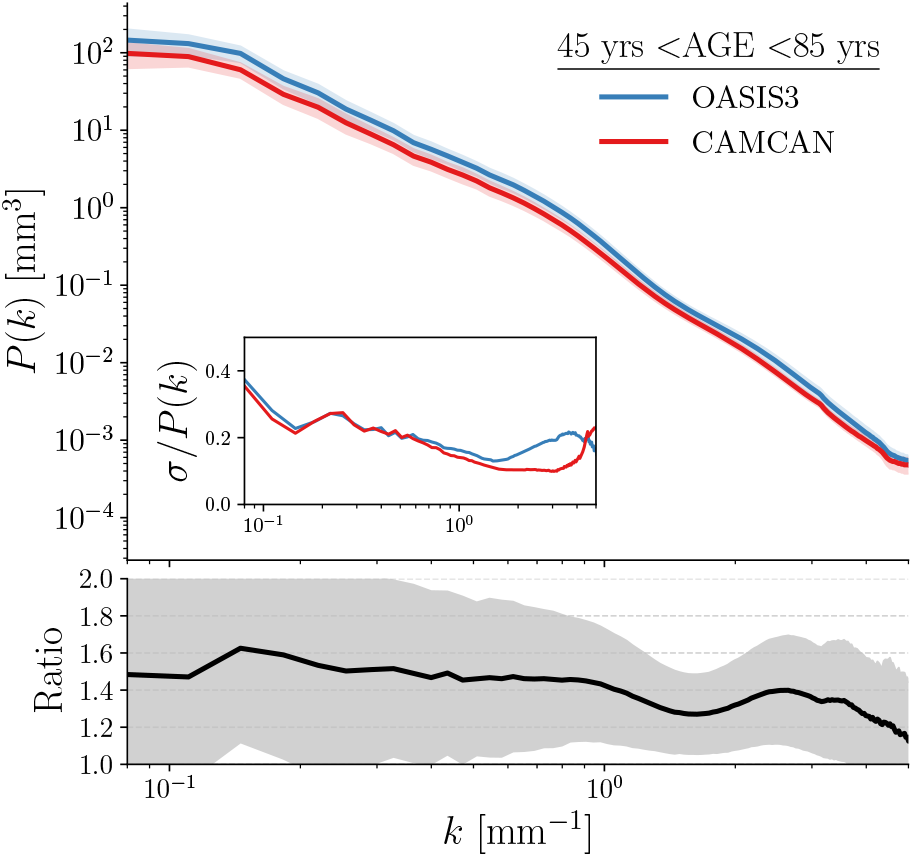
Comparison of the brain MRI power spectra for `OASIS-3` (blue) and `CamCAN` (red). Upper panel: power spectra *P* (*k*) as a function of spatial wavenumber *k*, shown on logarithmic axes. Both datasets exhibit a similar scale dependence, with `OASIS-3` displaying systematically higher power. Lower panel: the ratio *P*_OASIS-3_(*k*)*/P*_CamCAN_(*k*), highlighting a scale-dependent excess of approximately 10–60%. Shaded bands indicate 1*σ* standard deviation regions.

**Extended Data Fig. 3.**
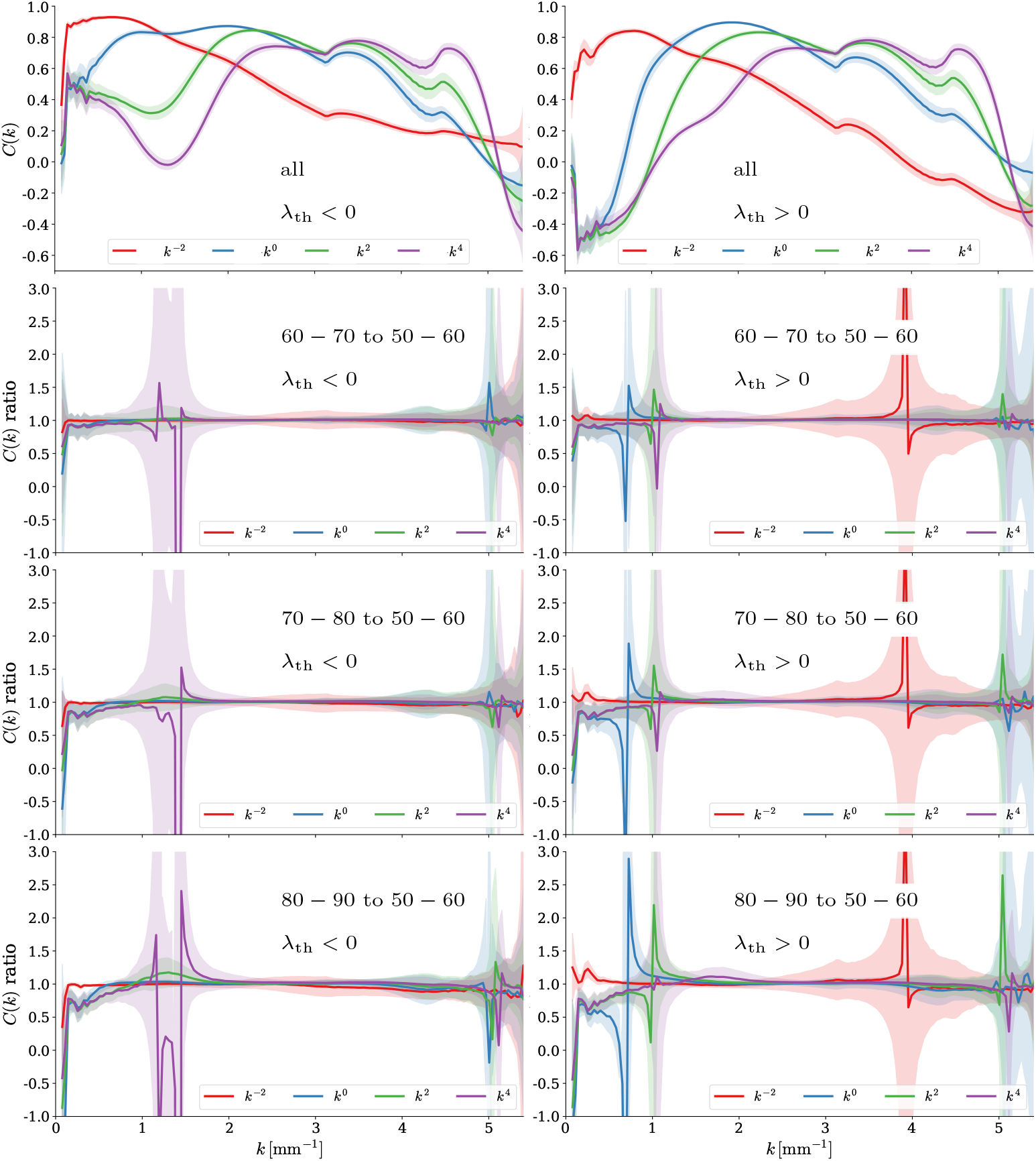
Image-field cross-correlations across age in OASIS-3. Panels compare the T1-weighted MRI intensity field with the voxel-wise threshold-averaged morphology fields at the four spectral levels, *s* ∈ {−2, 0, 2, 4}, with positive and negative threshold branches shown separately. The first row shows the cross-correlation curves for the age groups indicated; the panels below show the corresponding ratios to the reference curve specified in the plot. Narrow uncertainty bands and the close agreement of the age-group curves indicate a stable overall relationship between MRI intensity and the derived morphology, while the ratios highlight smaller relative differences.

**Extended Data Fig. 4.**
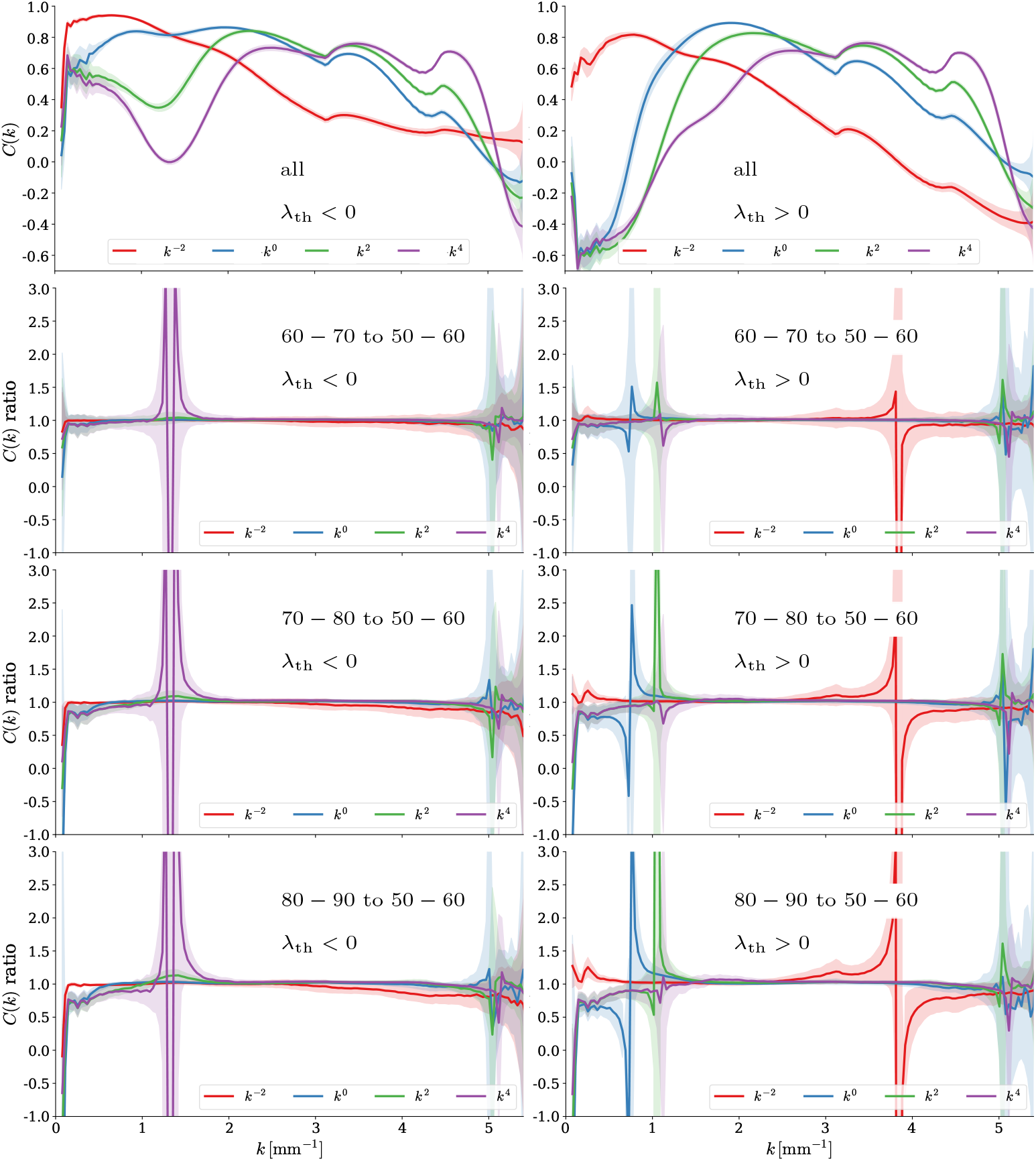
Image-field cross-correlations across age in CamCAN. Panels compare the T1-weighted MRI intensity field with the voxel-wise threshold-averaged morphology fields at the four spectral levels, *s* ∈ {−2, 0, 2, 4}, with positive and negative threshold branches shown separately. The first row shows the cross-correlation curves for the age groups indicated; the panels below show the corresponding ratios to the reference curve specified in the plot. Narrow uncertainty bands and the close agreement of the age-group curves indicate a stable overall relationship between MRI intensity and the derived morphology, while the ratios highlight smaller relative differences.

**Extended Data Fig. 5.**
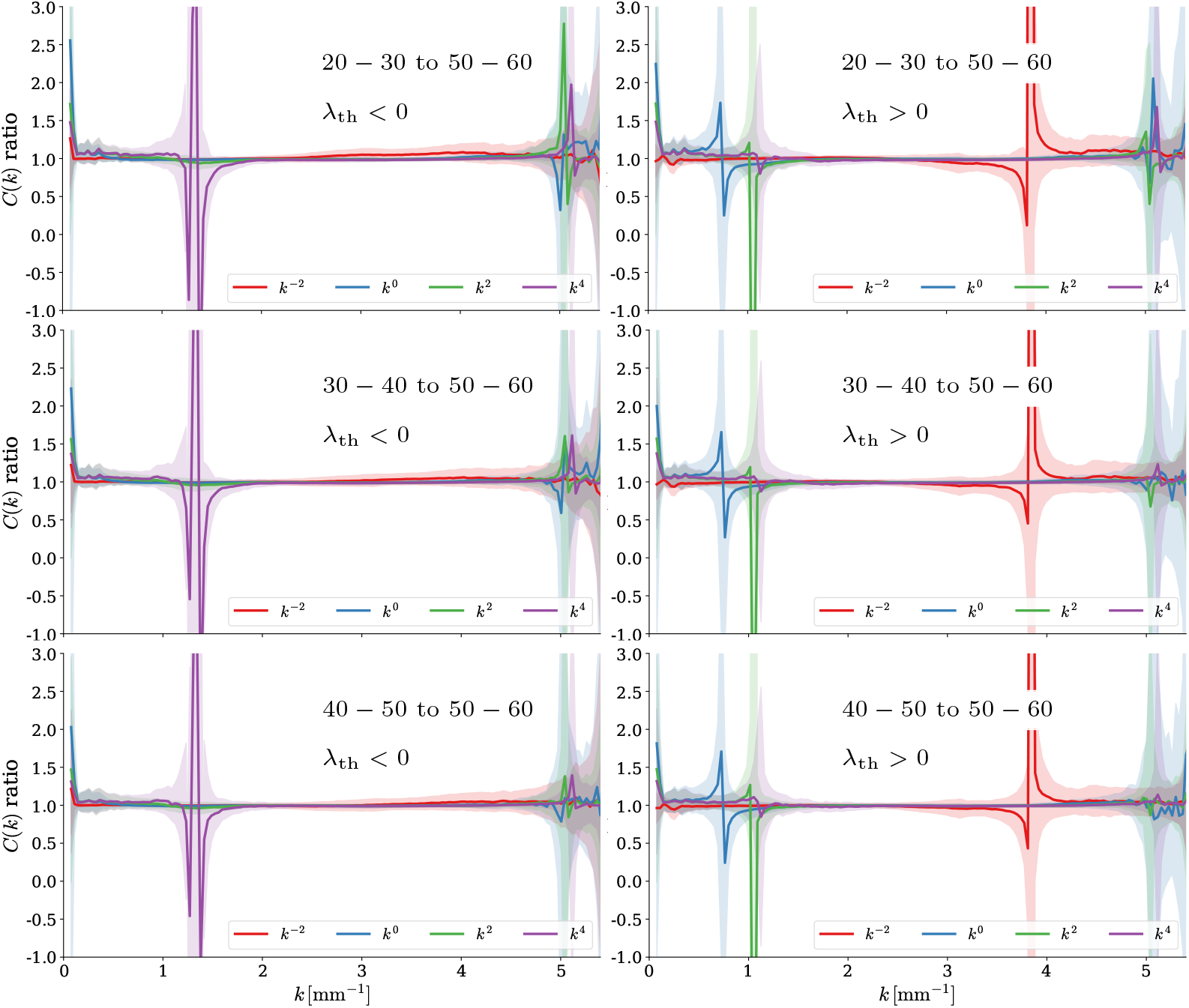
Additional age-group comparisons of image-field cross-correlations in CamCAN. Panels show cross-correlations between the T1-weighted MRI intensity field and the threshold-averaged morphology fields at *s* ∈ {−2, 0, 2, 4}, separately for positive and negative threshold branches. The first row shows the age-group curves, and the panels below show ratios relative to the reference specified in the plot, highlighting small differences between otherwise closely similar profiles.

**Extended Data Fig. 6.**
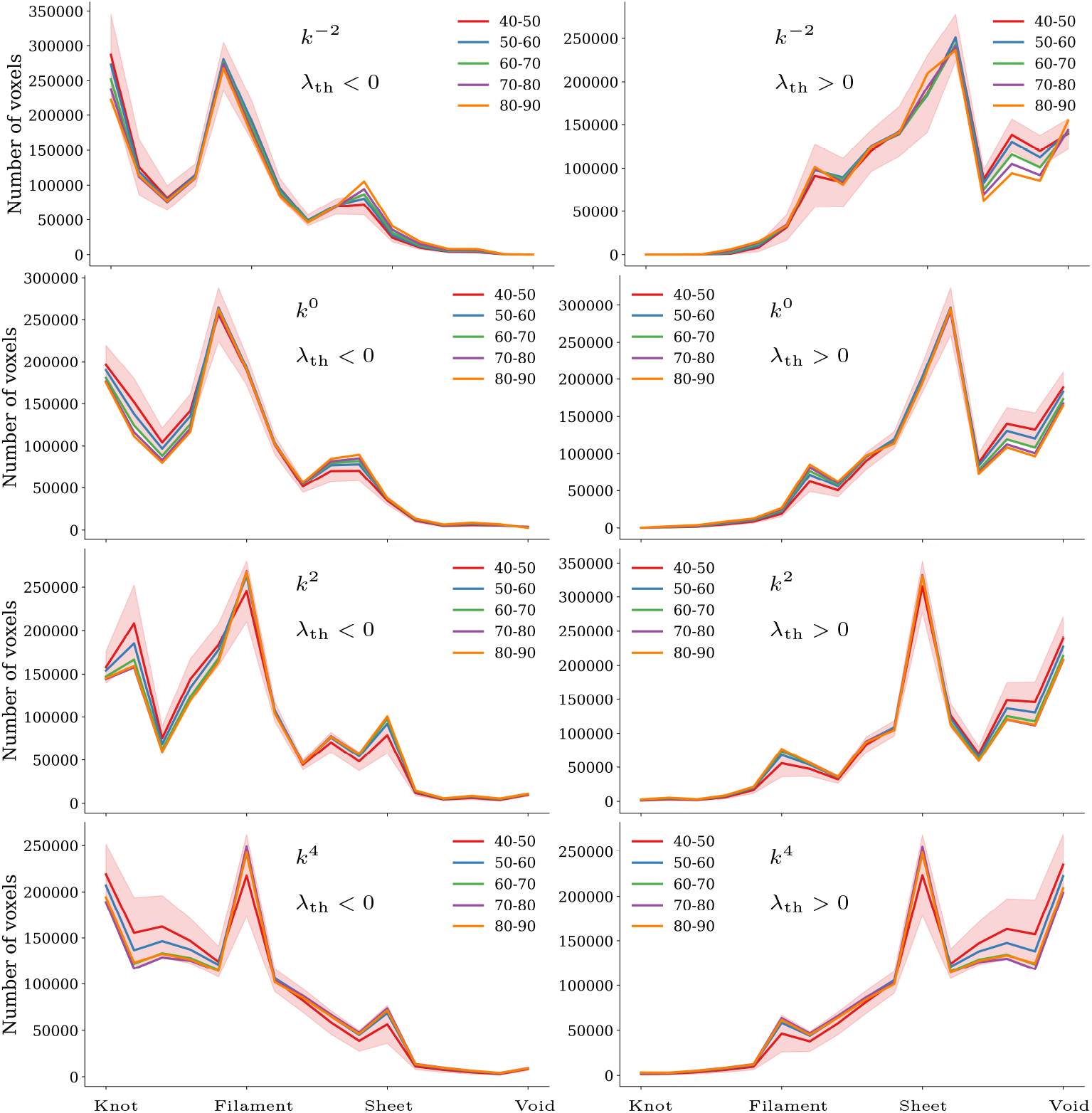
Threshold-averaged morphology across age in OASIS-3. Panels show histograms of voxel-wise classification labels averaged over eigenvalue thresholds at four spectral levels, *s* ∈ {−2, 0, 2, 4}. Positive and negative threshold branches are treated separately, with threshold magnitudes sampled up to the point at which the classification maps lose discernible morphological contrast.

**Extended Data Fig. 7.**
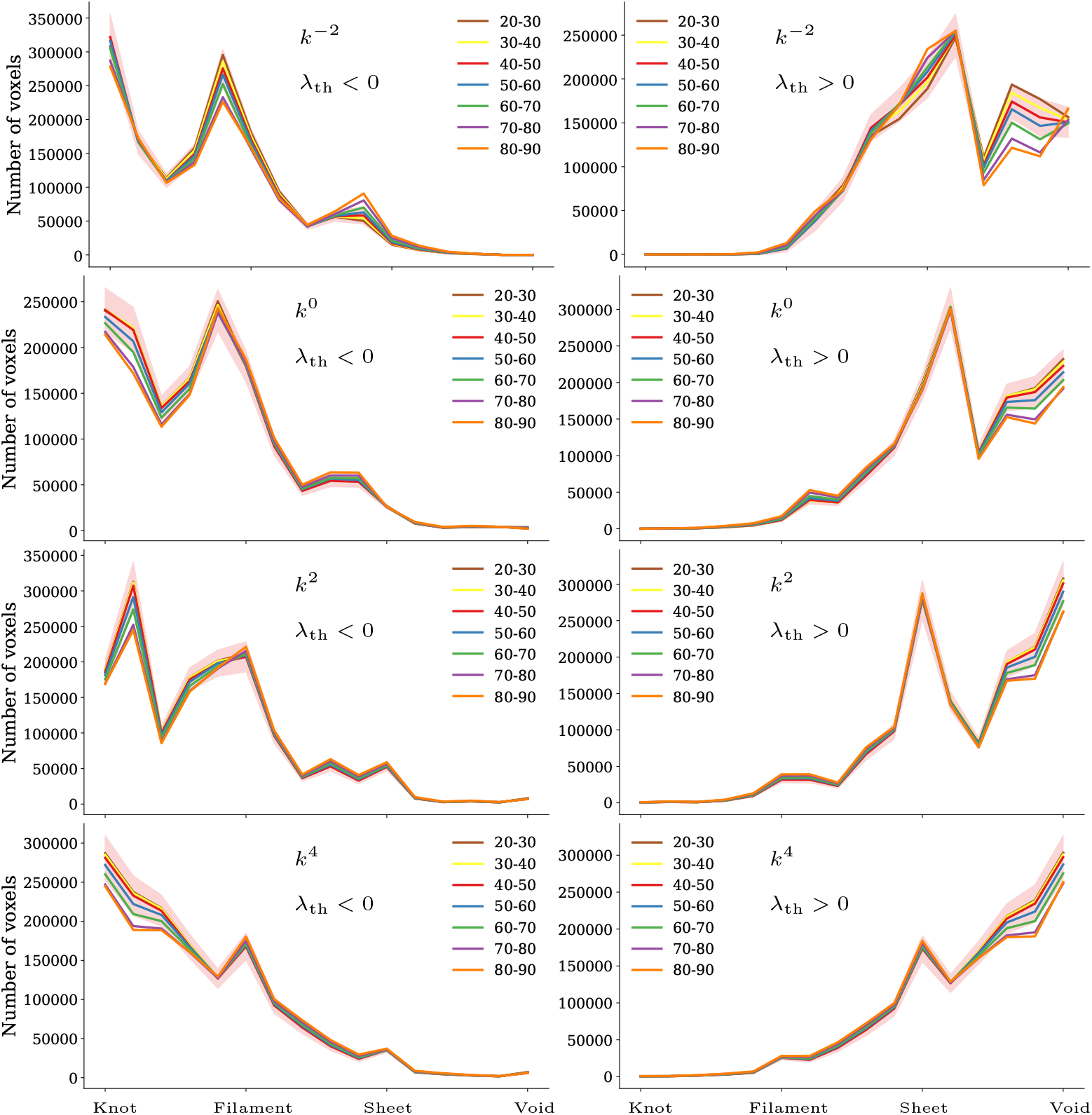
Threshold-averaged morphology across age in CamCAN. Panels show the four spectral levels, *s* ∈ {−2, 0, 2, 4}, each with separate positive and negative branches as in Extended Data Fig. 6.

**Extended Data Fig. 8.**
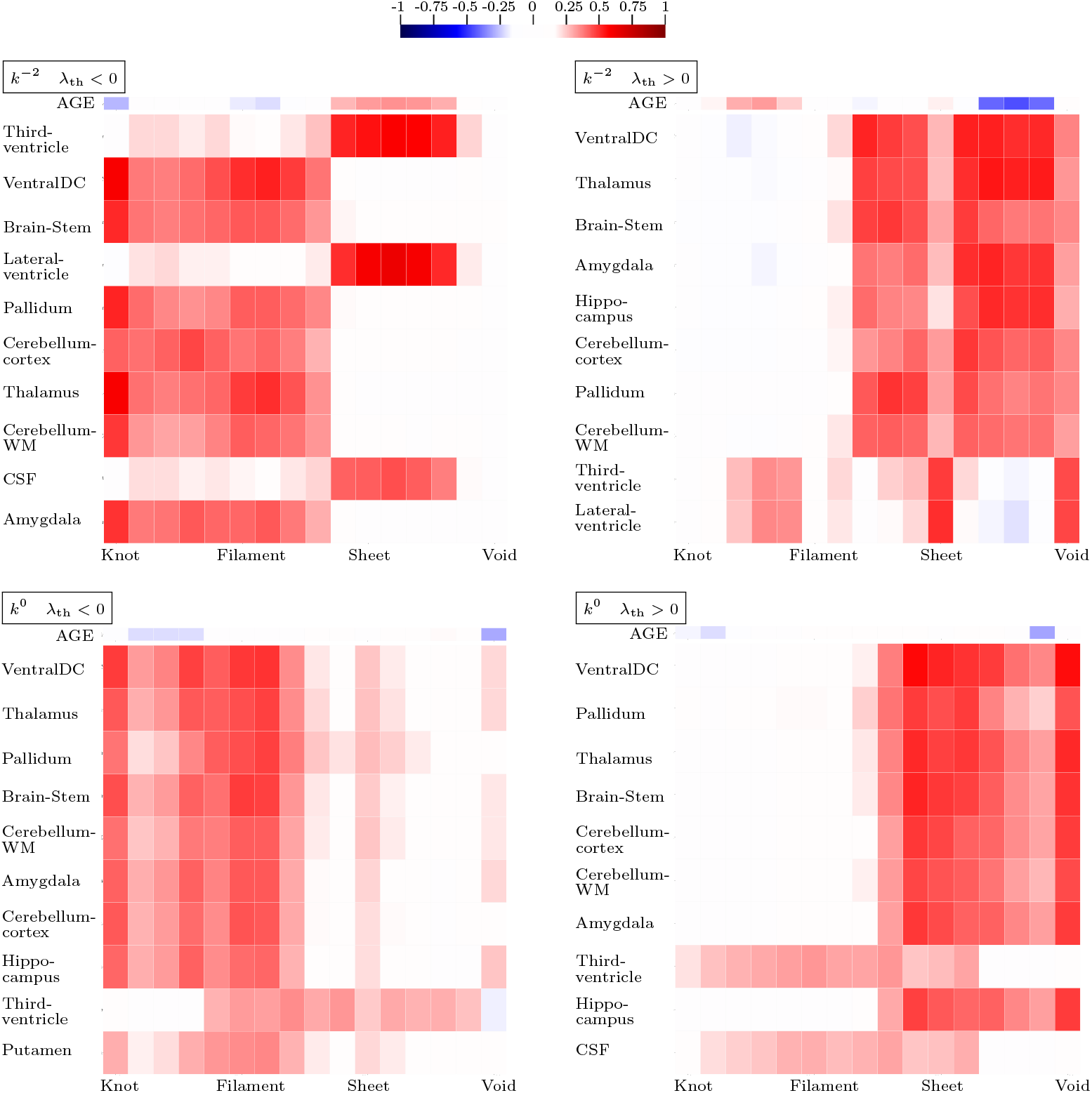
Associations of BWEB morphology with age and regional anatomy in OASIS-3. The top and bottom rows show the spectral levels *s* = −2 and *s* = 0, respectively, with negative eigenvalue-threshold branches (*λ*_th_ *<* 0) on the left and positive branches (*λ*_th_ *>* 0) on the right. Columns represent bins of the threshold-averaged morphology-score histograms, ordered from knots through filaments and sheets to voids. The upper strip shows Kendall’s rank correlation with chronological age; the heatmap below shows correlations with ten selected anatomical measurements derived from FreeSurfer-compatible segmentations. Red and blue indicate positive and negative correlations, respectively, with colour intensity indicating their magnitude. These across-subject associations differ from the spatial image-field cross-correlations in Extended Data Figs. 3–5.

**Extended Data Fig. 9.**
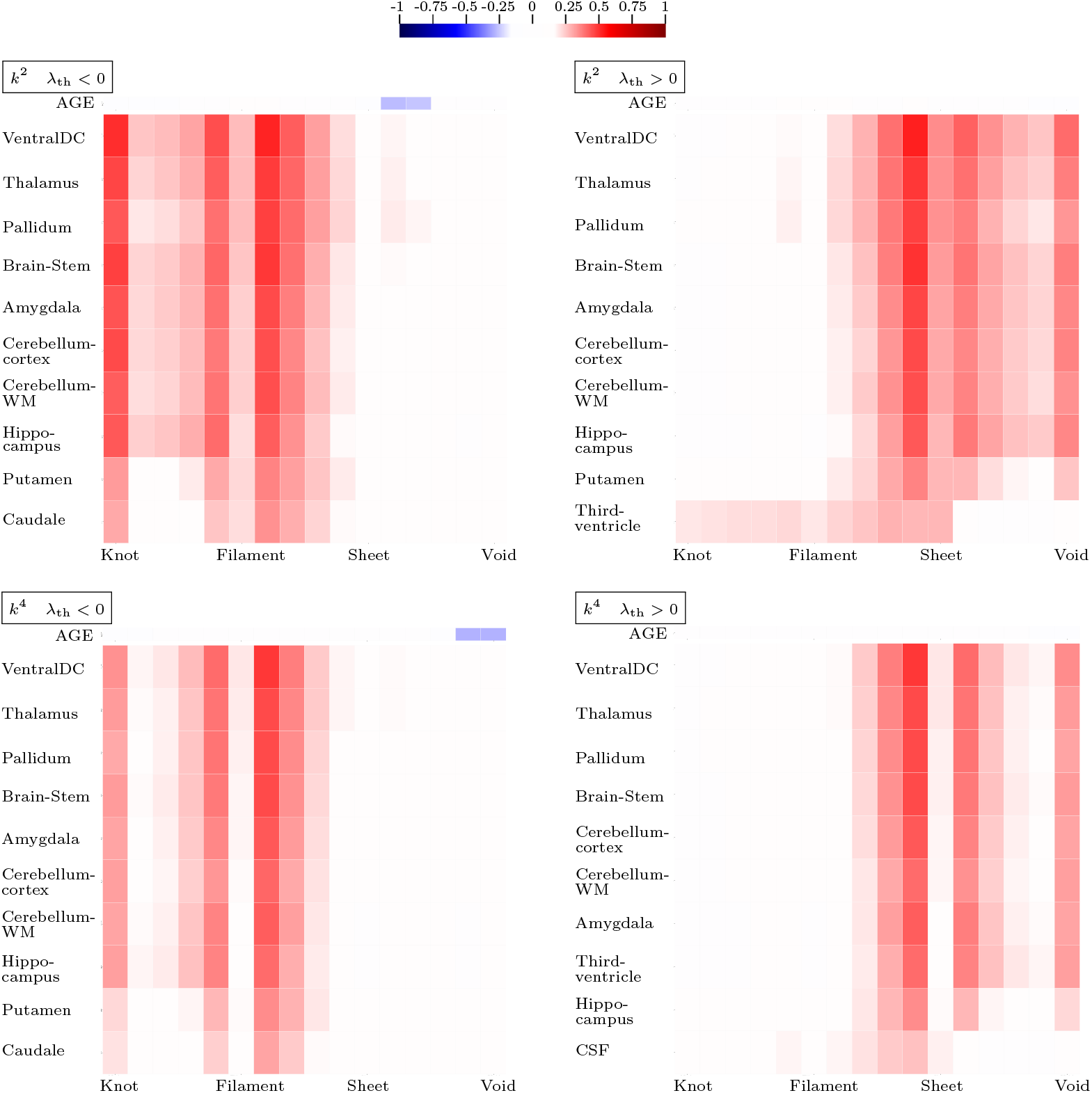
Same as Extended Data Fig. 8 but for levels *s* = 2 and *s* = 4.

**Extended Data Fig. 10.**
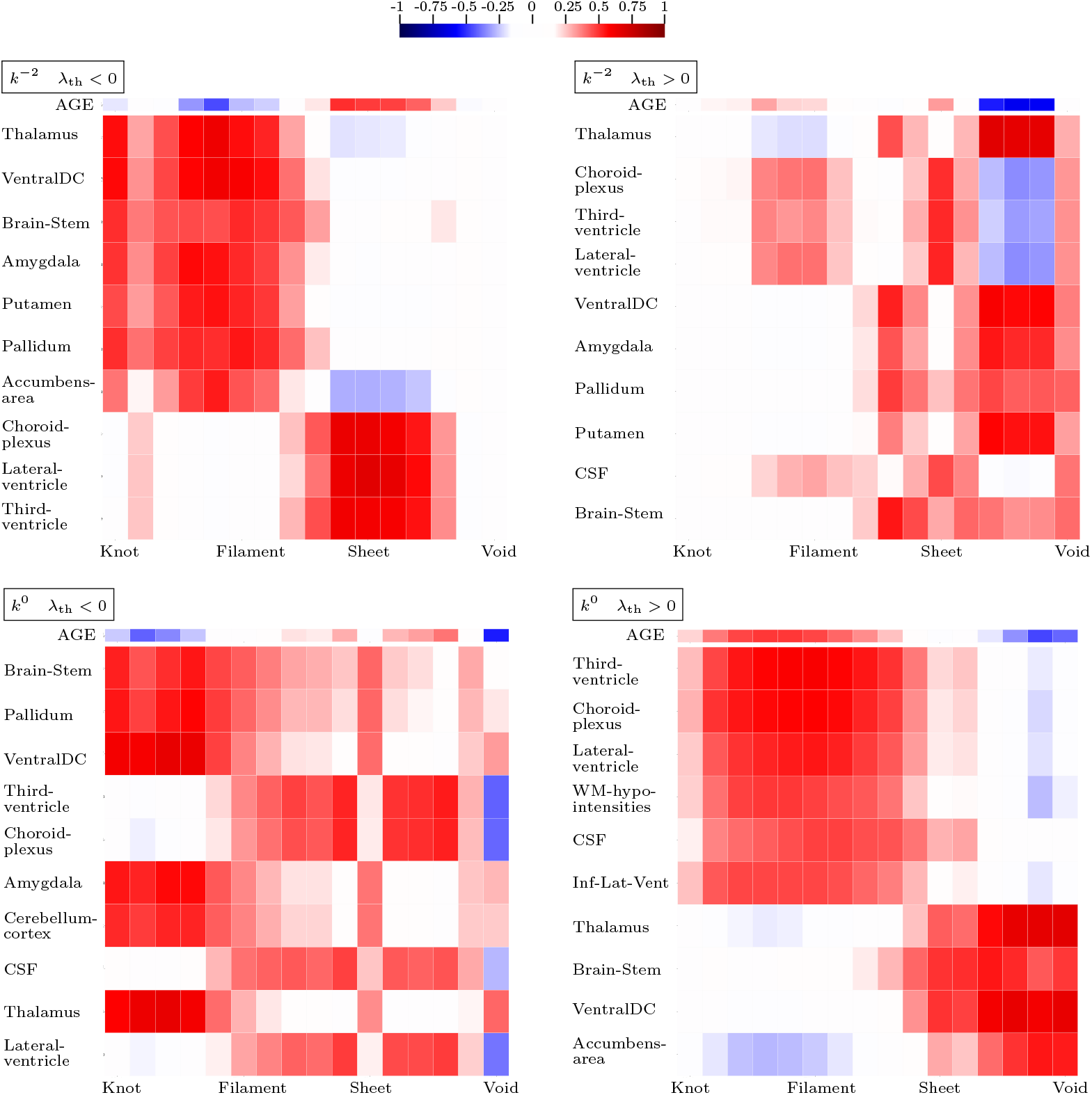
Same as Extended Data Fig. 8 but for `CamCAN`.

**Extended Data Fig. 11.**
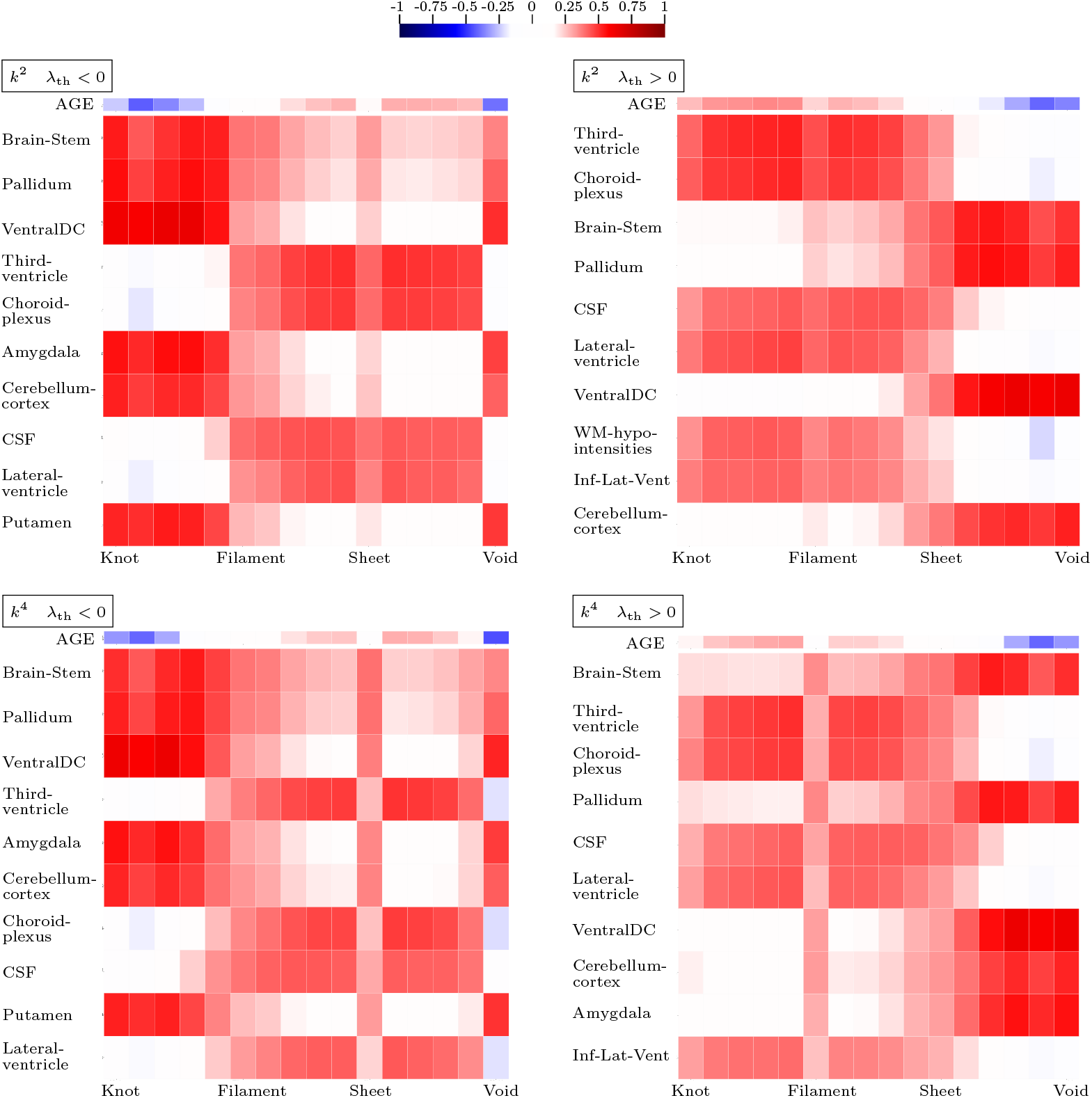
Same as Extended Data Fig. 10 but for levels *s* = 2 and *s* = 4.

## Footnotes

1 `SHARP` was developed within the COSMIC BRAIN research project (www.cosmic-brain.org).

